# Septo-hypothalamic regulation of binge-like alcohol consumption by the nociceptin system

**DOI:** 10.1101/2024.03.14.585116

**Authors:** Harold Haun, Raul Hernandez, Luzi Yan, Meghan Flanigan, Olivia Hon, Sophia Lee, Hernán Méndez, Alison Roland, Lisa Taxier, Thomas Kash

## Abstract

High intensity alcohol drinking during binge episodes overwhelmingly contributes to the socioeconomic burden created by Alcohol Use Disorders (AUD). Novel interventions are needed to add to the current therapeutic toolkit and nociceptin receptor (NOP) antagonists have shown promise in reducing heavy drinking days in patients with an AUD. However, an endogenous locus of nociceptin peptide and discrete sites of NOP action underlying this effect remains understudied. Here we show that the lateral septum (LS), a region contributing to binge drinking, is enriched in neurons expressing mRNA coding for the nociceptin peptide (*Pnoc)*. Pnoc-expressing neurons of the LS (LS^Pnoc^) are tuned to stimuli associated with negative valence and display increased excitability during withdrawal from binge-like alcohol drinking. LS^Pnoc^ activation was found to have aversive qualities and also potentiates binge-like drinking behavior, suggesting a convergence of circuitry that promotes aversion and drives alcohol consumption. Viral mediated tracing and functional assessment of LS^Pnoc^ projection fields revealed GABAergic synapses locally within the LS, and downstream within the lateral hypothalamus (LH) and supramammillary nucleus (SuM). Genetic deletion of NOP from the LS attenuated binge-like alcohol intake in male mice while NOP deletion from the LH and SuM decrease alcohol intake in females. Together, these findings are the first to demonstrate an endogenous population of nociceptin-expressing neurons that contributes to alcohol consumption and identifies sex-dependent modulation of alcohol drinking by NOP.

## INTRODUCTION

Excessive alcohol consumption is highly prevalent in the US and contributes to the ever-increasing socioeconomic burden produced by Alcohol Use Disorders (AUDs) ^1–3^. Binge drinking is the most common pattern of excessive drinking behavior and is shared across the population, in individuals diagnosed with an AUD as well as those that may drink excessively on occasion but do not meet the DSM criteria for an AUD^4–7^. Binge drinking is defined as the rapid consumption of alcohol containing beverages resulting in blood alcohol concentrations (BACs) in excess of the legal limit of intoxication (0.08 g/dl), typically consisting of 5+ standard drinks in men and 4+ for women in a session^8–10^. This risky pattern of drinking is the leading cause of emergency room visits in the US and correlated with a plethora of negative outcomes, such as organ damage, fatal automobile incidents, domestic and sexual violence, accidental injury, loss of workplace productivity, and is a strong predictor of the development of an AUD^5,6,11,12^. Presently, there are three FDA approved medications for the treatment of AUD, however, no singular “silver bullet” has been identified given that there is a high prevalence of relapse and resistance to pharmacological intervention^13–16^. Thus, there is a pressing need to expand our therapeutic toolkit to meet current demands.

Clinical and preclinical studies have identified the Nociceptin/Orphanin-FQ (nociceptin) peptide system as a candidate for AUD treatment development. In fact, a recent study demonstrated efficacy of a nociception receptor (NOP) antagonist to reduce heavy drinking days, and increase days abstinent, in treatment-seeking patients with an AUD^17^. Preclinical studies support this finding in that selective NOP antagonists attenuate alcohol self-administration and stress-induced reinstatement of alcohol seeking behavior in alcohol-preferring rats^18–20^, and reduce binge-like alcohol consumption in mice^21^. The uniformity of response to NOP antagonists to decrease alcohol drinking across preclinical models and rodent species is highly encouraging, warranting in-depth neuroanatomical study to isolate potential sites of action. Interrogating endogenous nociceptin-NOP circuitry is critical to understanding how the system may be dysregulated by alcohol and better inform treatment strategies. However, no studies to date have isolated and examined a nociceptin-expressing population in a preclinical model of AUD, driving the question of endogenous nociceptin involvement in excessive drinking behavior. The lateral septum (LS) is a site of interest given that it is enriched in *Pnoc* mRNA^22,23^, promotes drug seeking behavior^24–26^, and drives binge-like alcohol drinking in male mice^27^. Furthermore, the LS is situated as an integral node in addiction circuitry, interfacing directly with regions involved in binge drinking behavior such as the VTA, nucleus accumbens (NAc), and hypothalamus^28–30^. Thus, nociceptin-containing neurons in the LS (LS^Pnoc^) could be a previously unidentified node in circuitry mediating excessive drinking behavior.

The first goal of the present experiments was to classify expression patterns of nociceptin-coding mRNA (*Pnoc*) and NOP (*Oprl1*) throughout the neuroanatomical subdivisions of the LS. We then sought to determine activity patterns of LS^Pnoc^ during alcohol drinking, and general appetitive and aversive behaviors with fiber photometry. LS^Pnoc^ sensitivity to a history of binge-like alcohol consumption was assessed using electrophysiology and the reinforcing properties of LS^Pnoc^ activation was explored leveraging on optogenetic technique^31^. A chemogenetic approach^32^ was then used to bidirectionally modulate LS^Pnoc^ to determine a causal role in binge drinking behavior. Lastly, we employed channelrhodopsin-assisted circuit mapping to determine functional LS^Pnoc^ projection sites and used a genetic deletion strategy to determine the contribution of NOP in these downstream projection fields to binge drinking behavior.

## MATERIALS AND METHODS

### Subjects

Male and female Pnoc-IRES-cre and NOP*^fl/fl^* mice from our established breeding colony were housed in a temperature and humidity controlled AAALAC approved facility. Tail snips were collected from all mice and samples genotyped by Transnetyx prior to use. For all experiments, mice were singly housed at 8-10 weeks of age, maintained on a reverse 12-hr light/dark cycle, and provided with food (Isopro, RMH3000) and water *ad libitum*. Sex as a biological variable was defined in terms of chromosomal and gonadal sex as male or female for all studies. Pnoc-cre mice (N= 153) were used for fiber photometry, electrophysiology, chemogenetic, optogenetic experiments, and circuit mapping studies. A total of N= 22 were excluded from final analysis due to lack of viral expression and/or missed fiber placement. *In situ* hybridization and genetic deletion studies used N= 118 NOP*^fl/fl^* mice and 7 were excluded for lack of viral expression.

### Surgical Procedures

Mice underwent intracranial stereotactic surgery at 8-12 weeks of age using procedures previously described^27^. Briefly, mice were anesthetized with isoflurane (2%) and a Hamilton Neuros syringe was used to infuse adeno-associated virus (AAV) into either the LS, LH, or SuM at a flow rate of 0.05 uL/min, followed by a 5 min diffusion period, and 5 min retraction. For fiber photometry, electrophysiology, chemogenetics, optogenetics, and circuit mapping, mice received bilateral infusion (0.3 uL/side) of AAV constructs into the LS (AP: +0.8, ML: +/- 0.3, DV: −3.8), including: AAV8-hSyn-DIO-GCaMP7f (Addgene: 104488); AAV8-hSyn-DIO-mCherry (Addgene: 50459), AAV8-hSyn-DIO-hM4Di-mCherry (Addgene: 44362), AAV8-hSyn-DIO-hM3Dq-mCherry (Addgene: 44361), AAV8-EF1a-hSyn-DIO-synaptophysin-mCherry (MIT Vector core), or AAV8-EF1a-DIO-ChR2-mCherry (Addgene: 20297). Fibers were positioned above the LS (AP: 0.6; ML: 0.5; DV: −2.7) for photometry and optogenetic experiments and secured to the skull using Metabond as previously described^33^. For NOP deletion experiments, mice received bilateral infusion (0.3 uL/side) of AAV8-hSyn-cre-GFP (UNC Vector Core) or AAV8-hSyn-GFP (UNC Vector Core) into the LS (AP: +0.8, ML: +/- 0.3, DV: −3.8), LH (AP: −1.0, ML: +/-1.2, DV: −5.55), or a unilateral infusion into the SuM (5 degree angle; AP: −2.8, ML: +/-0.3, DV: −5.1). All mice were given at least 1 week of recovery after surgery prior to habitation for behavioral testing, and all experiments were conducted no sooner than 4 weeks after surgery to ensure sufficient viral expression.

### In Situ Hybridization (ISH)

Fresh frozen tissue was collected from NOP*fl/fl* mice for identification of cell type within the LS and confirm genetic deletion of *Oprl1* mRNA from the LS, LH, and SuM. Mice were placed in isoflurane induction chamber until deeply anesthetized, the brain was removed and immediately frozen on dry ice, then stored at −80c until sectioning. Serial slices from each region (14 uM) were collected on a cryostat (Leica CM3050 S) and mounted on Superfrost Plus slides. Slides were then processed for ISH using Advanced Cell Diagnostics RNAscope Fluorescence Multiplex Assay Kit according to the provided protocol. Briefly, mRNA was visualized with TSA Vivid Dyes (AKOYA Biosciences) targeting the vesicular GABA transporter (vGAT; Slc32a1 Probe# 319191), prepronocicpetin (Pnoc; Probe# 437881), and the nociceptin receptor (Oprl1; Probe#514301). Slides were coverslipped with Prolong Diamond mounting medium and imaged on VX200 Slide scanner (Olympus Life Sciences). Image analysis was conducted using QuPath software to determine *VGAT*, *Pnoc*, and *Oprl1* colocalization and a threshold was set at 1-5 puncta for positive expression within the footprint of DAPI labeling. Viral mediated NOP deletion was confirmed with ISH and mean *Oprl1* fluorescence intensity in each region was compared to control using QuPath software.

### Drinking Procedures

Alcohol drinking consisted of two paradigms that model binge-like alcohol consumption. Alcohol was prepared weekly and diluted in tap water from a 95% stock to a final concentration of 20% (v/v). Electrophysiology and tissue collection studies used the well-established “drinking in the dark” (DID) procedure^34,35^, consisting of 2-hr limited-access to either alcohol or water, 3-hr into the dark cycle for 3 consecutive days. The 4^th^ day extended the drinking session to a 4-hr period, and mice remained undisturbed in their cage for 3 days before starting the cycle again. This was repeated for 3 consecutive weeks and electrophysiology occurred 24-hr after the final 4-hr drinking session. Chemogenetic and genetic deletion studies used a modified DID procedure consisting of 2-hr limited access to alcohol, 3-hr into the dark cycle, for 5 consecutive days^27^. This modification was made to accommodate within-subjects testing during the 3^rd^ cycle of DID drinking, and produces blood alcohol concentrations (BACs) in excess of 80 mg/dl (**Supplemental Fig 3G, 4G, 5G**). Testing of water drinking occurred over a 24-hr period and sucrose (1%; w/v) drinking was tested over 2 weeks (Mon-Fri) in the 2-hr limited-access DID model described above. All fluid measurements were recorded immediately after the drinking session and adjusted for spillage.

### Fiber Photometry Hardware and Signal Processing

Neurophotometrics Inc equipment was used to record GCaMP7f-Ca transients from LS^Pnoc^ during various behaviors as previously described^36,33^. Briefly, hardware consisted of an LED driver box producing 470 and 415 nM light interleaved at 40 fps that was reflected through a dichroic mirror after bandpass filtering and collimation. LED light was focused through a 20x objective into a multibranch cord and output was measured and adjusted to ensure ∼50 uW output from the fiber tip for both 470 and 415 wavelengths. The emitted fluorescent signal was bandpass filtered and captured using a CCD camera at 40 fps. Bonsai software was used to trigger a USB camera to capture behavior aligned to the photometry signal, generate manual keydown timestamps, and Arduino-based TTL timestamps when necessary. Custom MATLAB code was used to deinterleave the 470 and 415 nm signals and subtract background florescence. The signal was fit to a bioexponential curve to correct for baseline drift and ΔF/F (%) was then generated (100*(signal-fitted signal)/(fitted signal)) for 470 and 415 signals. A z-scored signal was then calculated based on the entire recording session for the 470 output and fit using non-negative robust linear regression. The 415 output was then fit to the z-scored 470 output and subtracted for motion correction. The z-scored output for GCaMP7f was then aligned to timestamps generated for each behavior. The z-score for each behavioral test was plotted 5 s before and after bout or stimulus onset (with the exception of footshock given the prolonged decay time), and an average z-score was used for repeated behaviors or stimuli during a session (Ex: bouts of drinking). For analysis, the average z-score during the pre-bout interval was compared to the average post-bout z-score for all behavior/stimuli.

### Fiber Photometry Behavioral Testing

After surgery and recovery, mice were thoroughly habituated to patch cord tethering in Noldus Phenotyper boxes. Mice were connected to a dummy patch cord and allowed to explore the box for 30 min/day for at least 2 weeks prior to testing with food and water provided. For testing, mice were moved from the colony and sat undisturbed for no less than 1 hr prior to testing. A USB camera was used to record all behaviors and positioned for clear, unobstructed observation of behavior. Drinking experiments (water, sucrose, and alcohol consumption) were conducted during the dark cycle, under red light, in a modified home cage containing fresh cobb bedding with an open bonnet top. The home cage water bottle was removed prior to testing resulting in an acute 1-2 hr period of water deprivation for drinking experiments to encourage fluid consumption during testing. A single bottle containing alcohol, water, or sucrose was introduced after a 2 min acclimation period and timestamps were manually recorded for bouts of drinking. Bouts of drinking lasting at least 5 s were used for analysis and a final z-score was compiled as the average of 3 bouts (when 3 bouts were observed).

Splash testing was conducted in the modified home cage based on procedures from Quadir and colleagues^37^. After 2 min acclimation, mice were exposed to a spray of sucrose (10% w/v; in DI water), delivered from an atomizer (100 uL) to the dorsal coat. Mice were observed for 10 min and instances of grooming were recorded. Air puff testing was conducted in an open field arena measuring 50 x 50 cm. An air puff was applied 3 times to the dorsal coat for roughly 500 ms, and each exposure was separated by a 1 min interval. Elevated plus maze (EPM) testing was conducted under ambient room light. Mice were placed in the center of the EPM (75 cm) and center crossings, open arm entry, and open arm exit times were recorded for a 10 min period. Events were analyzed when time spent in the open arm was greater than 5 s. Footshock testing was conducted in a med associates chamber equipped with a shock grid flood. Footshock (0.6 mA for 2 s) was administered after a 3 min equilibration period, and this was repeated for a total of 5 footshock presentations separated by 90 s inter-shock interval.

### Electrophysiology

Tissue for electrophysiology experiments was rapidly extracted from anaesthetized (isolflurane) mice, sectioned at 250 uM on a vibratome (Leica VT100 S) in ice cold oxygenated sucrose (194 mM, NaCl: 20 mM, KCl: 4.4 mM, CaCl_2_: 2 mM, MgCl_2_: 1 mM, NaH_2_PO_4_: 1.2 mM, glucose: 10 mM, and NaHCO_3_: 26 mM), and transferred into oxygenated aCSF (NaCl: 124 mM, KCl: 4.4 mM, NaH_2_PO_4_: 1 mM, MgSO_4_: 1.2 mM, D-glucose: 10 mM, CaCl_2_: 2 mM, and NaHCO_3_: 26 mM) in a heated water bath (32-35 °C). After a 30-min equilibration period, tissue was transferred to a submerged recording chamber (Warner Instruments) under constant perfusion of heated aCSF (30 °C) at a flow rate of 2 mL/min. Cells were visualized with an Olympus BX51WI microscope and mCherry-expressing LS^Pnoc^ cells were identified with a CoolLED pE-100 (Andover, UK) optical stimulator. Thin walled borosilicate glass pipettes were pulled (P-97, Sutter Instruments) to 3–4 MΩ and filled with filtered, ice cold, K-gluconate (135 mM, NaCl: 5 mM, MgCl2: 2 mM, HEPES: 10 mM, EGTA: 0.6 mM, Na2ATP: 4 mM, Na2GTP: 0.4 mM, pH 7.3, 289–292 mOsm) or CsCl (cesium methanesulfonate: 134 mM, KCl: 10 mM, MgCl2: 1 mM, EGTA: 0.2 mM, MgATP: 4 mM, Na2GTP: 0.3 mM, phosphocreatine: 20 mM, pH 7.3, 285–290 mOsm with 1mg/mL QX-314). Data acquisition, digitization (10 kHz), and filtering (3 kHz) occurred through an Axon Multiclamp 700B amplifier (Molecular Devices). Analysis was the conducted in pClamp (Molecular Devices) and EasyElectrophysiology and cells were discarded in the event that a change in Ra exceeded 20% of the initial value.

For excitability experiments, recordings from fluorescent cells (LS^Pnoc^) began in voltage-clamp mode after a 5 min equilibration period, with membrane potential held at −70 mV with K-gluconate internal. Capacitance (Cm), input resistance (Rm), and access resistance (Ra) were recorded and excitability experiments were conducted in current-clamp mode. After a 2 min equilibration period with no holding current, resting membrane potential (RMP) and spontaneous firing activity was recorded from a 2 min gap-free recording (1 min epoch analyzed from 30s-90s). Current was then applied to hold the membrane potential at −70 mV (cells were excluded if the current necessary to hold at −70 mV exceeded 100 pA), and rheobase and voltage-current (V-I) plot protocols were run. The rheobase protocol consisted of ramping 100 pA steps and the V-I plot protocol consisted of 20 pA current steps (200 ms) starting at −100 pA and ending at 280 pA. Lastly, cells were returned to voltage-clamp mode and Cm, Rm, and Ra were recorded.

For ChR2-assisted circuit mapping experiments, tissue slices containing the LS, LH, or SuM were collected from mice expressing AAV8-hSyn-DIO-ChR2-mCherry in LS^Pnoc^. Recordings were conducted in voltage-clamp mode with CsCl internal in both mCherry+ and mCherry-neurons. After a 5 min equilibration period, blue light pulses (100 ms) were introduced at 1, 5, and 20 hz in 5 sweeps. Oxygenated aCSF containing picrotoxin (500 uM) was then applied to the tissue under continuous perfusion for 30 min. Sweeps of blue light stimulation were applied at 1, 5, and 20 hz once every 5 min for the duration. Representative traces were selected under aCSF perfusion, and after 20 min of picrotoxin application.

### Optogenetics and Real Time Place Testing

Mice were habituated to daily handling and tethering to a dummy patch cord for no less than 1 week prior to testing. Tethering consisted of 10 min free exploration whilst tethered in a modified home cage. Testing was conducted in a custom 2-chamber box (25cm x 50cm) with identical sides. The box was cleaned and fresh bedding introduced between subjects. Overhead video was collected and mice were tracked in real time using Ethovision software. TTL output was directed to a custom Arduino to generate a 20 hz frequency (2 ms pulse width) from a 473 nm laser (Shanghai Laser, ADR-800A). Patch cords were routed through a fiber optic rotary joint (Doric; FRJ 1×1 FC-FC) and laser output (∼5 mW) was checked from fiber tips prior to testing. Experimental testing consisted of a Pre-Test and Test phase. Pre-Testing was a 15 min free exploration session and time spent on either side was used to counterbalance laser pairing. Laser-pairing occurred the following day during a 30 min Test phase and time spent on the laser-paired side (% total time) was the main dependent variable.

### Chemogenetics

Mice were habituated for 1 week prior to behavioral testing with daily intraperitoneal (ip.) injections of saline (0.9%; 20 mL/kg). Mice received daily saline injections 30 min prior to testing through each phase of the experiment (alcohol drinking, locomotor activity, and sucrose drinking). For testing, Clozapine-*N*-Oxide (CNO; 3 mg/kg; HelloBio) was prepared fresh daily and dissolved in saline. Mice were then given an acute injection of CNO or saline in a within-subjects, counterbalanced design, 30-min prior to testing. When necessary, mice were tested to following day in the event of a bottle spill or environmental disturbance in the testing room.

### Open Field

Open field testing occurred in a 60 cm wide × 60 cm long × 40 cm deep SuperFlex open field arena (Omnitech Electronics, Accuscan, Columbus, OH) with a centralized house light (∼500 lx) as previously described^27^. For chemogenetic experiments, mice received an injection of saline (ip.) 30-min prior to testing and were introduced to the open field arena during a 30-min session. After habituation, mice were administered CNO or saline in a counterbalanced, within-subjects design, 30-min prior to testing in the open field across 2 consecutive days. This strategy was chosen allow for within-subjects testing and account for novelty and order effects that occur with repeated testing in an open field. Genetic deletion experiments included habituation to the behavior room and mice were tested in a single 30-min open field session. Behavior was tracked through beam breaks and analyzed with Fusion v6.5 software. The center of the open field was defined as a 10 cm square in the center of the assay. Data were collected in 1 min bins for the 30-min testing session and distance traveled (cm) and time spent in the center of the arena (s) were analyzed.

### Immunohistochemistry

Tissue from each experiment (excluding *in situ*) was collected for immunohistochemistry as previously described^27^. Mice were deeply anaesthetized and perfused with cold PBS (15 mL) followed by paraformaldehyde (PFA, 4%; 15 mL). Tissue was extracted, post-fixed in PFA overnight and moved to PBS prior to sectioning. Tissue was sectioned on a vibratome (VT 1200s, Leica Biosystems) at 40 uM and sections stored in PBS with azide (0.02%). Slices for photometry, optogenetics, and genetic deletion studies were mounted directly on SuperFrost Plus slides, coverslipped with Prolong Diamond, and imaged on a Keyence microscope as the endogenous GFP or mCherry tag was readily visible. Immunohistochemisty was used to amplify the mCherry tag for chemogenetic and synaptophysin tracing studies. Briefly, tissue was permeabilized in Triton X-100 (0.5%, 30 min) in PBS and blocked with normal donkey serum (NDS; 10%; 1 hr). Tissue was then incubated overnight at room temperature on a rocker in the primary antibody (mouse anti-RFP, 1:500; Rockland) in NDS. After washing the following day, tissue was incubated in the Alexa-Fluor 594 secondary antibody (1:200; donkey anti-mouse; Jackson Immuno) in 0.1% Triton X-100 for 2 h. Tissue was then mounted on SuperFrost Plus slides and coverslipped with Prolong Diamond mounting media. Images were collected on a Keyence microscope at 4x, 10x, and 20x magnification.

### Statistical Analysis

Data were analyzed by ANOVA or t-test, significance threshold set to 0.05, and significant main effects and factor interactions are reported in **Supplementary Table 1**. Image analysis for *in situ* hybridization studies included sex (male, female) as a between subjects variable, and AP gradient (anterior, intermediate, posterior), and DV gradient (dLS, iLS, vLS) were within subjects factors for each probe. Fiber photometry experiments included sex as a between subjects variable, time (pre, post bout average z-score) was the within subjects factor, and z-score was the main dependent variable. For electrophysiology experiments, sex and drinking history (water, alcohol) were between subjects variables, and current was a within-subjects factor. The dependent variables were input resistance (Ohm), capacitance (pF), resting membrane potential (mV), number of action potentials in a 1 min period, rheobase (pA), and action potentials per step. Optogenetics included sex and AAV (ChR2, mCherry) as between subjects factors and day (PreTest, Test) as a within subjects variable. Chemogenetic studies included sex and AAV (hM4Di, hM3Dq, and mCherry) as between subjects factors, and drug treatment (vehicle, CNO) as a within subjects variable. Genetic deletion studies included sex and AAV (Cre, GFP) as the main factors, and time (Week 1, 2, 3) as a within subjects comparison. Dependent variables for chemogenetic and genetic deletion studies included alcohol intake (g/kg), sucrose intake (mL/kg), water intake (mL/kg), body weight (g), distance traveled (cm), and time (s) spent in the center of an open field. *Post hoc* analysis were planned comparisons as a function of bout (pre, post) for photometry experiments, treatment (water, alcohol) for electrophysiology experiments, day (PreTest, Test) for optogenetics, drug (vehicle, CNO) for chemogenetic experiments, and AAV (Cre, GFP) for genetic deletion studies. All *post hoc* comparisons used Sidak’s correction for planned multiple comparisons. Analyses were conducted using SPSS software (v24) and GraphPad Prism 9 (La Jolla, CA). Data are presented as mean +/- standard error.

## RESULTS

### The Lateral Septum is Enriched in Pnoc and Oprl1 mRNA

The LS is a GABAergic basal forebrain structure comprised of dorsal (dLS), intermediate (iLS), and ventral (vLS) subdivisions along the dorsal-ventral (DV) gradient, and spans roughly 4 mM along the anterior-posterior (AP) gradient in mice^38–40^. Because nociceptin and NOP expression has not been rigorously canvased within the LS, we used *in situ* hybridization to quantify *VGAT*, *Pnoc,* and *Oprl1* mRNA in NOP*fl/fl* mice (**Fig. 1A-E**). Quantification was conducted in DAPI+ cells within the 3 subdivisions of the LS (dLS, iLS, and vLS) across the AP gradient (**Fig. 1B**). For analysis, male and female data were collapsed across sex given a lack of main effect or factor interaction.

**Fig. 1:**
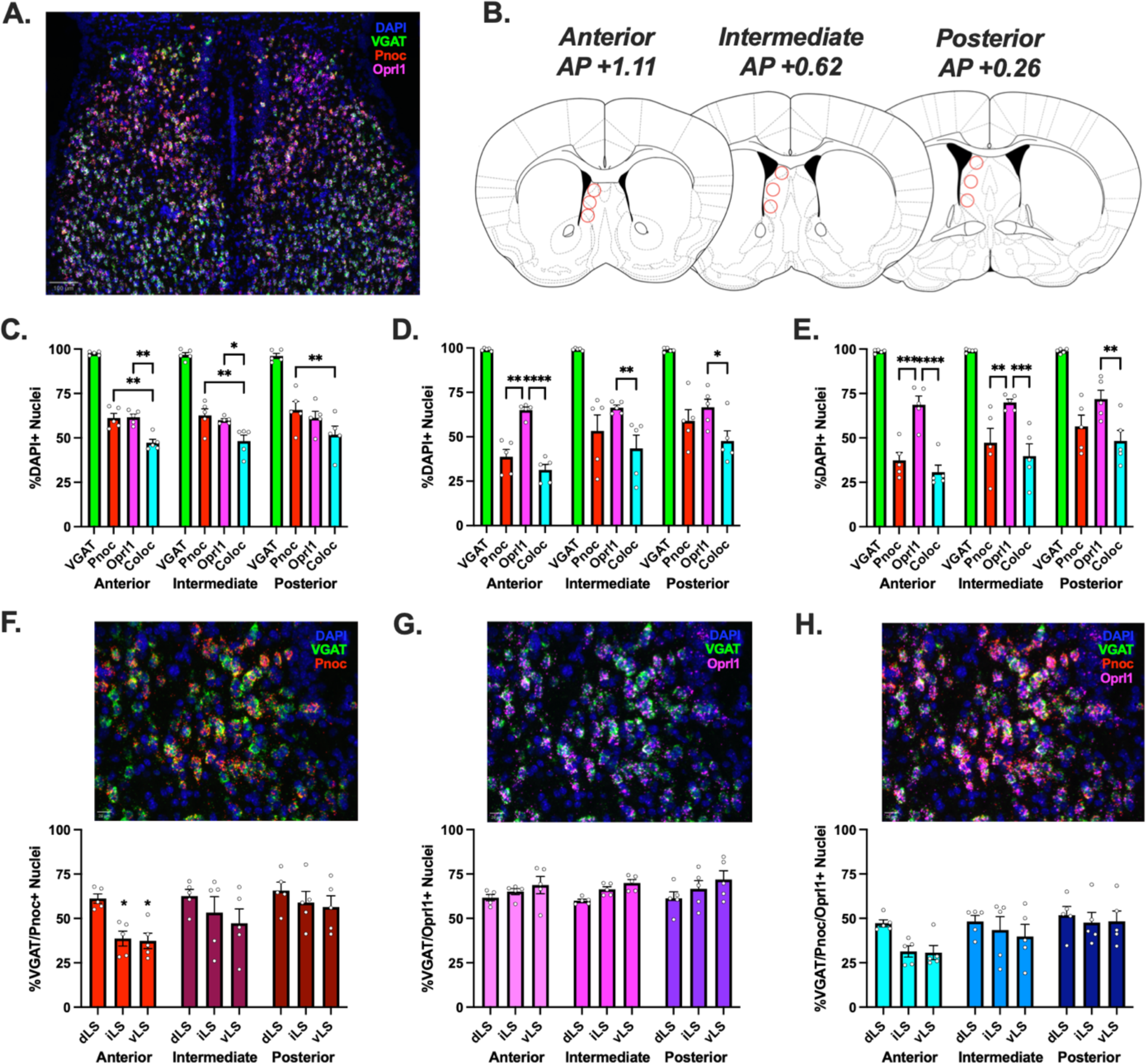
The Lateral Septum is Enriched in *Pnoc* and *Oprl1* mRNA. (A.) Representative image of the LS visualizing probes for *VGAT* (GABA), *Pnoc* (Nociceptin), and *Oprl1* (NOP) mRNA in DAPI+ nuclei at 5x magnification. Scale bar = 100 uM. (B.) Representation of ROIs for analysis from the anterior, intermediate, and posterior dLS, iLS, and vLS. (C.-E.) Quantification of mRNA within the dLS, iLS, and vLS. Values are collapsed across sex given the lack of main effect and factor interaction. (C.) Roughly 60% of cells within the anterior dLS express *Pnoc* or *Oprl1* mRNA, and *Pnoc/Oprl1* colocalization is less that *Pnoc* and *Oprl1* expressing cells alone. (D.) *Oprl1* is highly enriched in the iLS and expression is greater than *Pnoc* in the anterior subdivision and *Pnoc/Oprl1* colocalization in all subdivisions. (E.) *Pnoc* is expressed to a lesser extent than *Oprl1* in both anterior and intermediate vLS. (F.) Representative 20x image of *VGAT* and *Pnoc* expression in the LS. Pnoc is more highly expressed in the anterior dLS compared to iLS and vLS, and the posterior dLS is enriched in *Pnoc* compared to anterior iLS and vLS. Scale bar = 20 uM. (G.) Representative 20x image of *VGAT* and *Oprl1* expression in the LS. Roughly 65% of cells within the LS express *Oprl1* mRNA, with no differences between subregion. (H.) Representative 20x image of *VGAT*, *Pnoc*, and *Oprl1* expression in the LS. Roughly 45% of cells within the LS express both *Pnoc* and *Oprl1* mRNA. Data are represented as mean +/- SEM (*P< 0.05, **P< 0.01, ***P< 0.005, ****P< 0.001).

As expected, roughly 98% of cells within the LS expressed *VGAT* mRNA (**Fig. 1C-E**). *Pnoc* expression was most abundant within the dLS (63%; **Fig. 1C**) and to a lesser extent in the anterior iLS (39%; **Fig. 1D**) and vLS (37%; **Fig. 1E**). In contrast, *Oprl1* expression (66%) was fairly consistent across the AP and DV gradients of the LS suggesting a ubiquitous pattern of expression. *Pnoc* and *Oprl1* colocalized across the LS within 43% of cells with no significant differences within the AP and DV subdivisions. This indicates the possibility of feedback regulation of nociceptin and/or GABA release locally within the LS, as this structure is reported to be highly interconnected^39,41,42^. Together, these data suggest that *Pnoc* expressing neurons are most dense within the dLS independent of AP position and *Oprl1* is uniformly expressed.

### LS^Pnoc^ Respond to Stimuli Associated with Negative Valence

An abundance of Pnoc-expressing neurons were observed in the LS and this region as a whole is reported to be involved in consummatory behavior as well as aversion-related behaviors^29,30^. A series of fiber photometry experiments were conducted to selectively assess LS^Pnoc^ responsivity to appetitive drinking behavior and exposure to aversive stimuli. AAV8-hSyn-DIO-GCaMP7f was infused into the LS of Pnoc-cre mice and fibers were positioned above this site to record calcium transients from LS^Pnoc^ (**Fig. 2**). After habituation to the tethering procedure, a single bottle containing tap water, alcohol (20%; v/v), or sucrose (1%; g/v) was introduced in separate tests and GCaMP activity was recorded and aligned with licking behavior. LS^Pnoc^ activity was increased preceding bouts of water drinking in male but not female mice (**Fig 2B**). While no change in activity was detected during bouts of alcohol drinking, LS^Pnoc^ activity decreased during active licking bouts for sucrose in males (**Fig. 2D**). This is consistent with activity recorded from LS-NT^43^ and -CRFR2^44^ which display decreased activity during consumption of a palatable reward. Alternatively, the discrepancy in signal between water, alcohol, and sucrose indicates precise tuning relative to the rewarding properties of these liquids. Because discrete cell types within the LS are tuned to aversive stimuli^44–46^, we then assessed LS^Pnoc^ activity in response to footshock and air puff exposure. Footshock elicited a robust increase in LS^Pnoc^ Ca activity in males and females (**Fig. 2E**), as did exposure to an acute air puff (**Fig. 2F**). A sucrose splash test was then conducted to query LS^Pnoc^ activity during grooming behavior, which is proposed to encode relief from the aversive state generated by a soiled coat. As with footshock and air puff exposure, an atomized spray of sucrose (10%; g/v; 100uL) to the dorsal coat resulted in increased LS^Pnoc^ activity (**Fig. 2G**). However, activity decreased during active bouts of grooming behavior post-splash exposure (**Fig. 2H**). Lastly, anxiety-like behavior was assessed in an EPM and elevated LS^Pnoc^ activity observed during exploration of the open arm was normalized upon return to the closed arm in males (**Fig. 2J**). Together, these findings indicate that LS^Pnoc^ activity increases during exposure to aversive stimuli and activity is decreased during relief-associated grooming behavior and during consumption of a palatable reward in males.

**Fig. 2:**
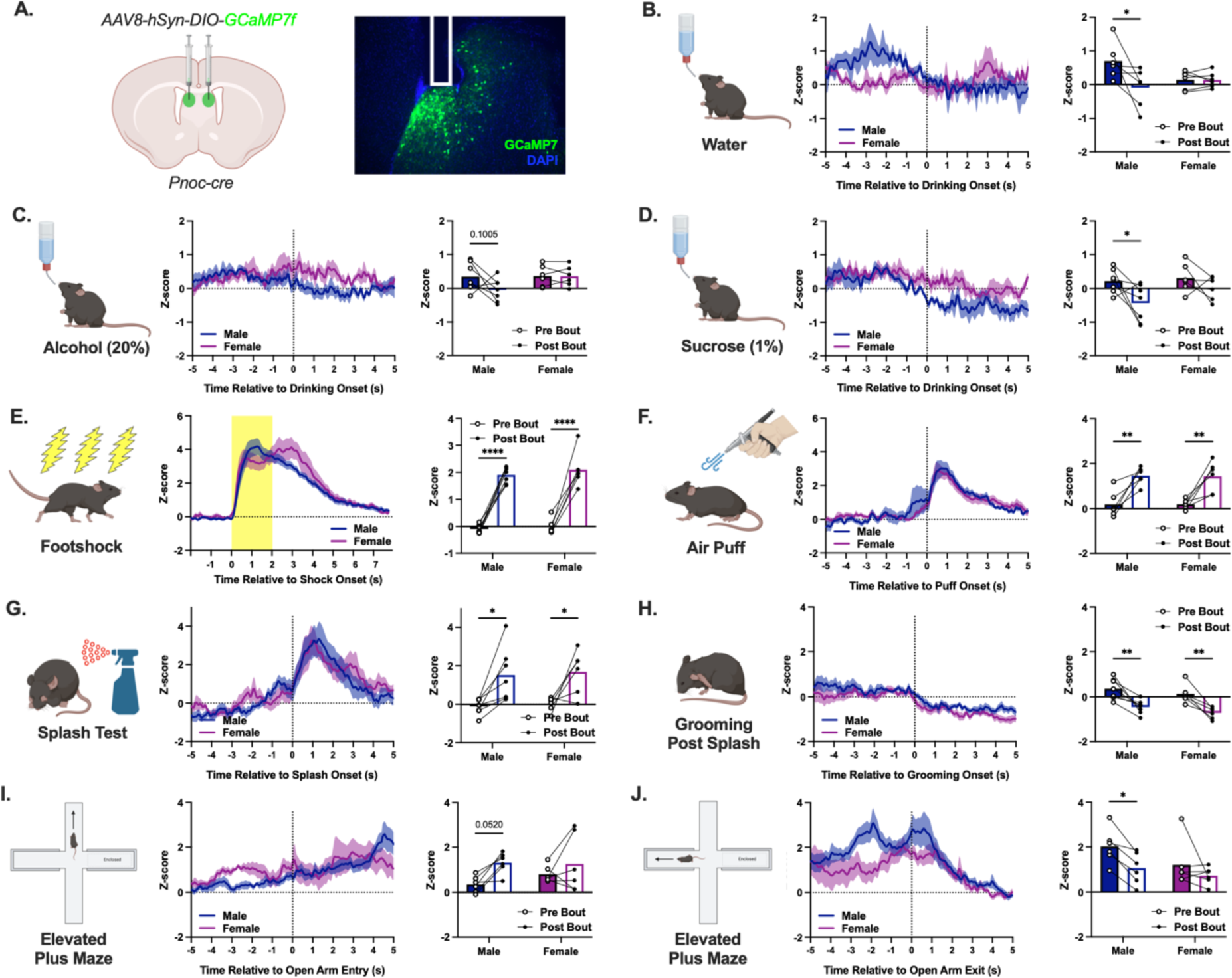
LS^Pnoc^ Respond to Stimuli Associated with Negative Valence. (A.) Schematic and representative image of viral infusion, GCAMP7f expression, and fiber placement within the LS. Full extent of viral footprint and fiber placements are shown in **Supplementary Fig. 1.** (B.) LS^Pnoc^ activity during bouts of water drinking was increased in male mice preceding licking onset (Bout *[F (1,10) = 5.90, P= 0.036]*). (C.) LS^Pnoc^ activity was not changed when mice were given access to alcohol. (D.) LS^Pnoc^ activity decreased in male mice during licking bouts for sucrose (Bout *[F (1,11) = 6.54, P= 0.027]*). (E.) Foot shock elicited robust LS^Pnoc^ activity in both male and female mice (Bout *[F (1,11) = 170.10, P< 0.001]*). (F.) Air puff applied to the dorsal coat resulted in elevated LS^Pnoc^ activity in males and females (Bout *[F (1,11) = 39.47, P< 0.001]*). (G.) Application of an atomized sucrose solution to the dorsal coat was associated with increased LS^Pnoc^ activity (Bout *[F (1,11) = 20.16, P< 0.001]*). (H.) Grooming behavior post sucrose splash was accompanied by a reduction in LS^Pnoc^ activity in both males and females (Bout *[F (1,11) = 39.87, P< 0.001]*). (I.) Mice were tested in the EPM and entry into the open arm resulted in a trend toward increased LS^Pnoc^ activity in males (Bout *[F (1,10) = 7.38, P= 0.022]*). (J.) Entry into the closed arm was associated with decreased LS^Pnoc^ activity in males when compared to the open arm (Bout *[F (1,10) = 9.19, P= 0.013]*). Data are represented as mean +/- SEM (*P< 0.05, **P< 0.01, ***P< 0.005, ****P< 0.001).

### Binge Drinking Increases Excitability of LS^Pnoc^

The LS is generally involved in binge-like alcohol consumption^27^ and alcohol withdrawal is associated with increased excitability in various brain regions and cell types associated with negative affect^47–51^. Because LS^Pnoc^ is responsive to aversive stimuli, we sought to assess LS^Pnoc^ excitability during withdrawal from binge drinking when signs of negative affect emerge using whole-cell patch-clamp electrophysiology. AAV8-hSyn-DIO-mCherry was infused into the LS of Pnoc-cre mice to selectively target LS^Pnoc^ and tissue was prepared for recording 24-hr after the final drinking session in mice with a history of water or alcohol drinking (**Fig. 3A**). Females generally consumed more alcohol than males and both consumed high levels of alcohol during the 4-hr drinking session (**Fig. 3B**)

**Fig. 3:**
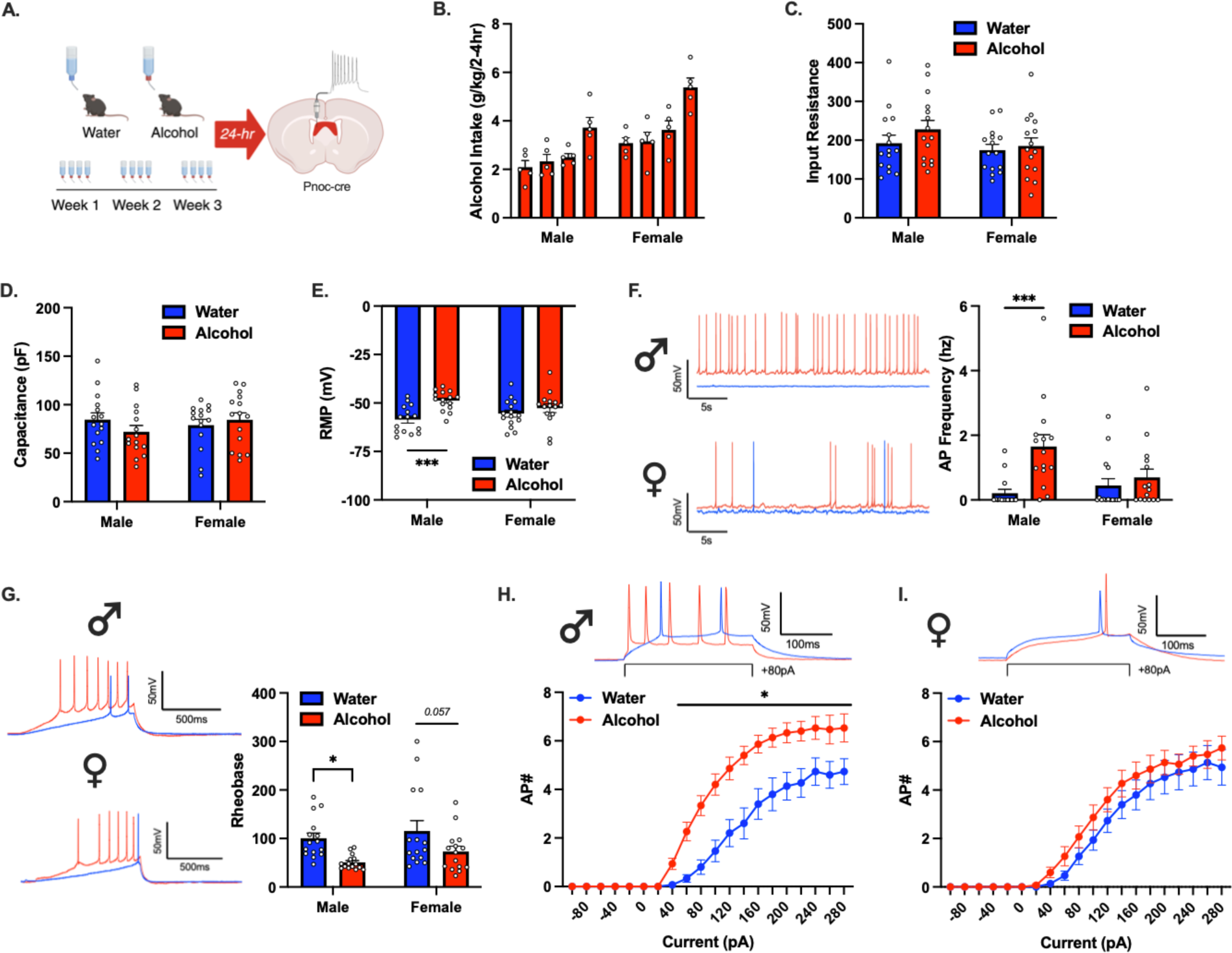
Binge Drinking Increases Excitability of LS^Pnoc^. (A.) Depiction of study design. Whole-cell patch-clamp electrophysiology was conducted 24-hr after the final binge drinking session and excitability was assessed in LS^Pnoc^. (B.) Average alcohol intake (g/kg) for males and females across 3 weeks of drinking the DID model. Females generally consumed more alcohol than males (Sex *[F (1, 8) = 10.16, P= 0.013]*) and drinking was greater during the 4-hr drinking session (Time *[F (3,24) = 30.19, P< 0.001]*). (C.-D.) Input was resistance and capacitance of LS^Pnoc^ was not altered as a function of drinking history in male and female mice. (E.) Male mice with a history of alcohol drinking displayed a depolarized shift in LS^Pnoc^ resting membrane potential (RMP) compared to water drinking controls (History *[F (1, 56) = 11.78, P= 0.001]*). (F.) LS^Pnoc^ action potential (AP) frequency (hz) was greater in males consuming alcohol than water (History *[F (1, 56) = 11.23, P= 0.001]*; Sex*History *[F (1, 56) = 5.583, P= 0.025]*). (G.) Rheobase was significantly decreased in LS^Pnoc^ as a function of alcohol drinking in males (History *[F (1, 56) = 11.88, P= 0.001]*). (H.) Current step protocols indicated increased firing at 40-180 pA in males with a history of alcohol drinking compared to water drinkers *(Current [F (1, 56) = 367.22, P< 0.001]*; History *[F (1, 56) = 10.89, P= 0.002]*; Current*History *[F (1, 56) = 5.70, P= 0.02]*). (I.) Alcohol drinking did not alter AP activity in female mice. Data are represented as mean +/- SEM (*P< 0.05, **P< 0.01, ***P< 0.005, ****P< 0.001).

Excitability was assessed in current-clamp mode through 4 metrics: resting membrane potential (RMP), spontaneous action potential (AP) frequency, rheobase, and APs generated through an increasing current-step protocol. Three weeks of binge-like alcohol consumption had no effect on input resistance or capacitance in LS^Pnoc^, but resulted in decreased RMP and increased AP frequency 24-hr post alcohol drinking compared to water-drinking control mice. The ability of alcohol drinking to influence LS^Pnoc^ RMP was driven in large part by male mice, as alcohol consumption resulted in a more depolarized RMP compared to water (**Fig. 3C**). Similarly, LS^Pnoc^ AP frequency was greater in male mice with a history of alcohol drinking compared to water, with no difference in females (**Fig. 3D**). Current was injected to hold cells at −70 mV and a current ramp protocol was used to assess rheobase, the minimum current required to elicit an AP. Drinking history resulted in a shift in rheobase with male mice in the alcohol drinking condition displaying a decrease compared to water drinkers, and this effect neared significance in females (**Fig. 3F**). Excitability assessed through a current step protocol revealed an increased number of AP events in male mice with a history of binge-like alcohol consumption with no difference in females (**Fig. 2G**; **Fig. 2H**).

### LS^Pnoc^ Activation is Aversive and Increases Alcohol Drinking

A major component of the current theoretical framework proposed for AUD involves excessive alcohol consumption to relieve the negative affective state driven by repeated cycles of binge intoxication and withdrawal, or drinking in the form of negative reinforcement^49,52^. To determine if LS^Pnoc^ are actively involved in the expression of aversion, a real-time place preference assay was conducted in a 2-chamber testing apparatus. The LS of male and female Pnoc-cre mice was infused with AAV-hSyn-DIO-ChR2/mCherry and fibers were positioned above this site (**Fig. 4A**). Fibers and viral footprint were largely confined to the dLS and iLS (**Supplemental Fig. 2A**). Mice were thoroughly habituated to handling and tethering prior to testing. Mice were then introduced to a 2-chamber apparatus with identical sides and time spent on either side of the chamber was recorded. During this pre-test session, mice spent equal time in either side of the 2-chamber box, and data were collapsed across sex given the lack of main effect or factor interaction (**Fig. 4C**). Side bias was controlled for based on the pre-test session and laser stimulation of LS^Pnoc^ in mCherry and ChR2-expressing groups was conducted the following day. ChR2-expressing mice spent less time on the laser paired side of the chamber, suggesting that LS^Pnoc^ activation resulted in avoidance behavior indicating aversive properties (**Fig. 4C**), and this effect was not observed in the control group (mCherry).

**Fig. 4:**
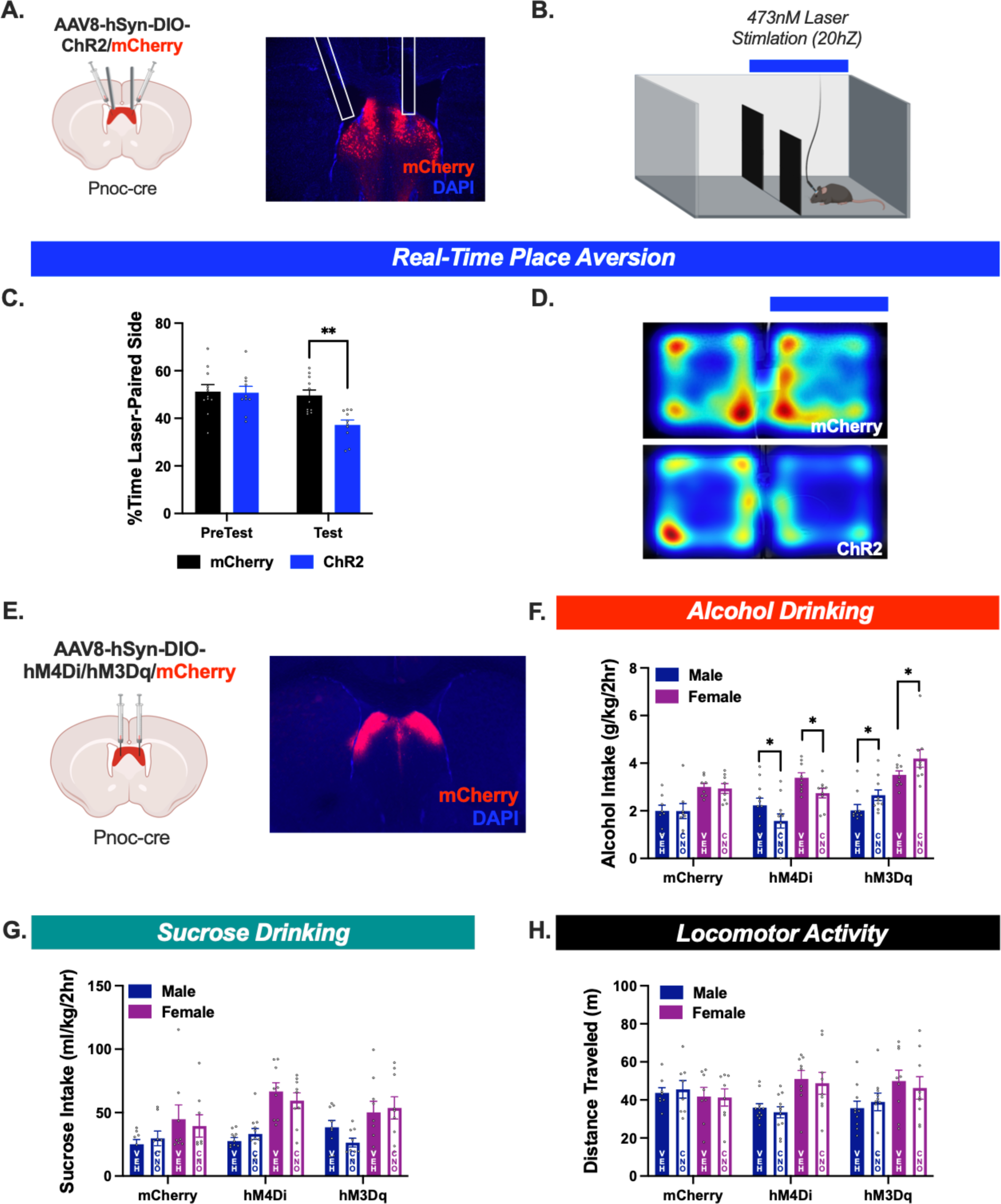
LS^Pnoc^ Activation is Aversive and Increases Alcohol Drinking. (A.) Depiction of viral infusion and representative image of mCherry expression and fiber placement in LS^Pnoc^ for optogenetic studies. Full extent of viral footprint and fiber placements are shown in **Supplementary Fig. 2A**. (B.) Diagram of 2 chamber testing apparatus with laser-paired side. (C.) While there was no difference in basal side preference during the PreTest phase, blue-light laser stimulation pairing resulted in decreased side preference in ChR2-expressing mice with no change in mCherry controls (Day*AAV *[F (1, 19) = 5.13, P= 0.035]*). Data are collapsed across sex given the lack of main effect. (D.) Representative heatmap of laser stimulation in mCherry- and ChR2-expressing mice. (E.) Depiction of viral infusion and representative image of mCherry expression in LS^Pnoc^ for chemogenetic studies. Heatmaps depicting viral spread can be found in **Supplementary Fig. 2B**. (F.) Mice were challenged with vehicle (VEH, saline) or CNO (3 mg/kg) in a within-subjects counterbalanced designed 30 min prior to alcohol drinking. CNO treatment in hM4Di-expressing male and female mice resulted decreased alcohol intake compared to vehicle. Male and female mice expressing hM3Dq in LS^Pnoc^ consumed greater levels of alcohol when challenged with CNO, suggesting that chemogenetic activation potentiated alcohol consumption. There was no effect of CNO on alcohol drinking in mCherry controls, but females consumed more alcohol than males in all groups (Sex *[F (1, 49) = 50.94, P< 0.001]*; AAV *[F (2, 49) = 5.86, P= 0.005]*; Drug*AAV *[F (2, 49) = 10.06, P< 0.001]*). (G.) There was no effect of LS^Pnoc^ chemogenetic manipulation on sucrose intake, although females consumed more sucrose than males (Sex *[F (1, 48) = 25.16, P< 0.001]*). (H.) There was no change in locomotor activity resulting from LS^Pnoc^ silencing or activation and females displayed a greater distance traveled than males (Sex *[F (1, 49) = 5.66, P= 0.021]*). Time spent in the center of the open field was not affected by CNO treatment across groups (**Supplementary Fig. 2C**). Data are represented as mean +/- SEM (*P< 0.05, **P< 0.01, ***P< 0.005, ****P< 0.001).

Because LS^Pnoc^ are highly active during withdrawal from binge drinking, and activation of this population has aversive properties, a chemogenetic experiment was conducted to determine the functional contribution of LS^Pnoc^ to alcohol drinking. We hypothesized that activation of LS^Pnoc^ drives a negative affective state that promotes alcohol drinking behavior, and that silencing of this population would decrease binge-like alcohol consumption. To test his hypothesis, AAV8-hSyn-DIO-hM4Di/hM3Dq/mCherry was infused into the LS of male and female Pnoc-cre mice (**Fig. 4E**), and viral expression was largely confined to the dLS and iLS (**Supplemental Fig. 2B**). Mice were challenged with vehicle (saline) and CNO (3 mg/kg) in a within-subjects counterbalanced design after habituation to alcohol drinking, sucrose drinking, and locomotor activity. Generally, females consumed more alcohol (20%; v/v) than males and CNO challenge was without effect in the control groups (mCherry). However, silencing of LS^Pnoc^ in hM4Di-expressing mice resulted in a reduction of alcohol intake in both males and females compared to vehicle (**Fig. 4F**). Conversely, activation of LS^Pnoc^ after CNO treatment in hM3Dq-expressing mice increased alcohol drinking in males and females. To determine if this effect was specific to alcohol, LS^Pnoc^ silencing/activation was then assessed in the context of sucrose (1%; w/v) drinking in an identical design (**Fig. 4G**). Sucrose was chosen as a contrast to alcohol given its reinforcing and caloric properties, and the concentration was selected as mice consume similar volumes of each solution within a 2-hr session. Females consumed more sucrose than males but there was no difference between vehicle and CNO treatment among AAV groups. Lastly, mice were assessed in an open field test to determine if the changes in alcohol intake could be attributed to a non-specific perturbation of locomotor activity. While females displayed greater overall locomotor activity, there was no effect LS^Pnoc^ manipulation on distance traveled (**Fig. 4H**) or time spent in the center of the open field (**Supplemental Fig. 2C**). Together, these findings indicate that LS^Pnoc^ are active during and contribute to aversive stimuli exposure, highly active during withdrawal from alcohol, and promote excessive drinking behavior.

### NOP Deletion from LS, LH, or SuM Attenuates Alcohol Intake

We next wanted to understand potential sites of action by which the peptide, nociception, may act via its receptor, NOP, to drive alcohol consumption. As a first step, we evaluated the projection patterns of LS^Pnoc^ neurons. Two complementary circuit mapping experiments were conducted to determine the projection patterns of LS^Pnoc^. AAVs harboring synaptophysin-mCherry or ChR2-mCherry were expressed in LS^Pnoc^ (**Fig. 5A**). Synaptophysin-mCherry expression was observed locally within the LS, indicating local innervation by LS^Pnoc^ (**Fig. 5B**). To confirm functionality, ChR2-evoked synaptic input onto non-Pnoc expressing neurons within the LS was tested using whole-cell patch-clamp electrophysiology. Blue light-evoked (1 hz) post-synaptic current was observed in non-Pnoc expressing neurons within the LS. This evoked current was blocked in the presence of picrotoxin (500 uM), suggesting that LS^Pnoc^ form local functional GABAergic synapses (**Fig. 5B**). Synaptophysin-mCherry immunofluorescence was also observed in the lateral hypothalamus (LH) (**Fig. 5C**). ChR2-assisted circuit mapping revealed optically evoked post-synaptic current within the LH. This evoked current was blocked by picrotoxin, confirming a functional GABAergic synapse within the LH arising from LS^Pnoc^ (**Fig. 5C**). Lastly, synaptophysin-mCherry immunofluorescence was observed in the supramammillary nucleus (SuM) of the hypothalamus (**Fig. 5D**). Blue light stimulation in LS^Pnoc^-ChR2 terminal fields within the SuM resulted in picrotoxin-sensitive evoked current, suggesting monosynaptic GABAergic connectivity (**Fig. 5D**).

**Fig. 5:**
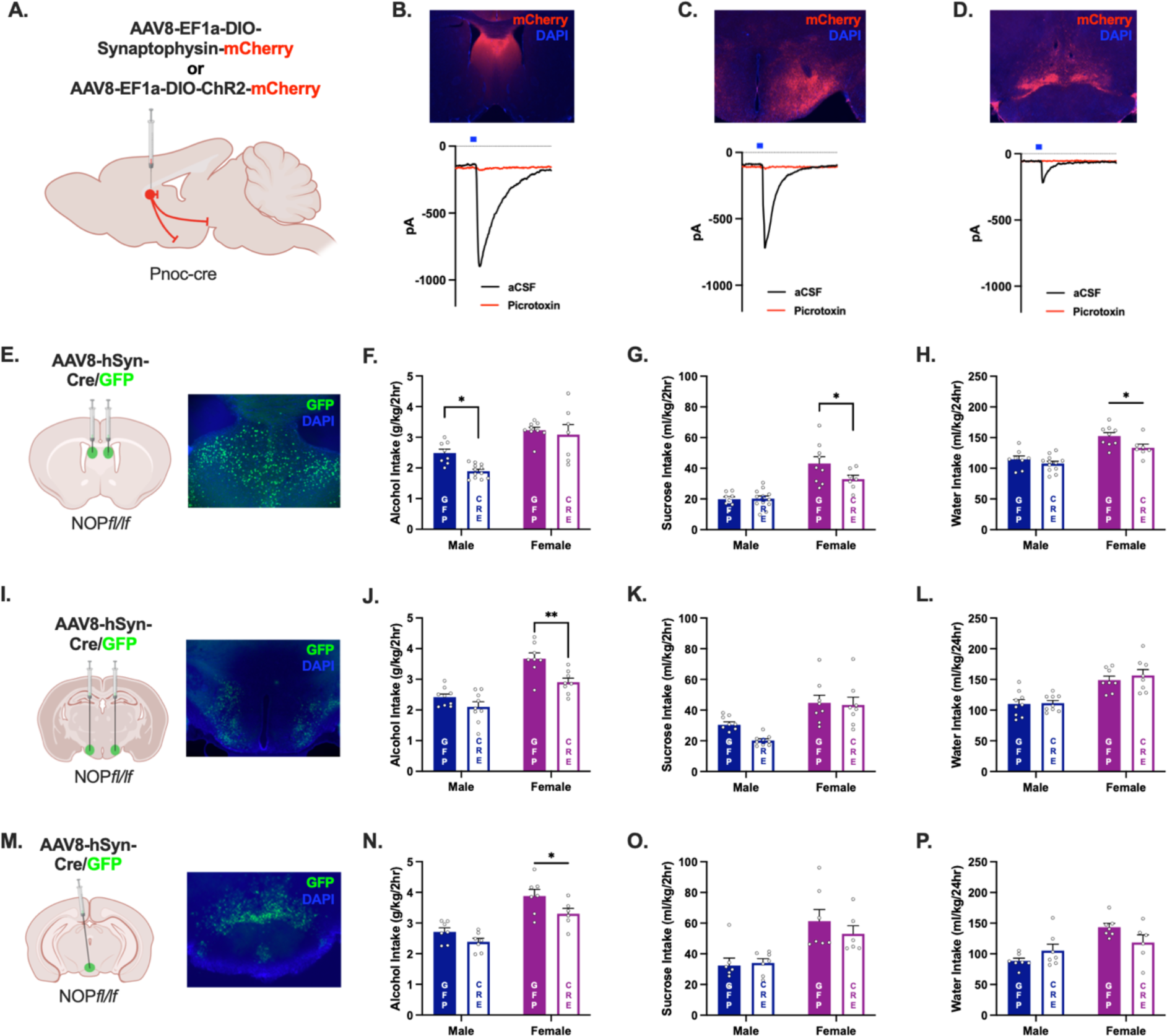
NOP Deletion from LS, LH, or SuM Attenuates Alcohol Intake. (A.) Depiction of viral infusion and tracing of LS^Pnoc^ in projection sites. (B.) Expression of synaptophysin-mCherry was observed in LS^Pnoc^ terminals locally within the LS. Stimulation of LS^Pnoc^ elicited an inhibitory post synaptic current (IPSC) in Pnoc-neurons within the LS. Picrotoxin bath application abolished the ChR2-evoked IPSC indicating a monosynaptic GABAergic circuit. (C.) LS^Pnoc^ terminal fields in lateral hypothalamus (LH). ChR2-evoked post synaptic current from LS^Pnoc^ afferents is blocked within the LH in the presence of picrotoxin. (D.) LS^Pnoc^ terminal fields in supramammillary nucleus (SuM). ChR2-evoked post synaptic current from LS^Pnoc^ afferents is blocked within the SuM in the presence of picrotoxin. (E.) Depiction of viral infusion in the LS and representative image of expression. Quantification of viral mediated NOP deletion and heatmaps visualizing viral spread within the LS are shown in **Supplemental Fig. 3A-D**. (F.) Genetic deletion of NOP from the LS in males resulted in decreased average alcohol intake compared the GFP-expressing controls (Sex *[F (1, 32) = 38.72, P< 0.001]*; AAV *[F (1, 32) = 5.61, P= 0.024])*. (G.-H.) Female mice expressing Cre in the LS consumed less sucrose than GFP in 2 hrs (Sex *[F (1, 32) = 18.26, P< 0.001]*), and less water over a 24 hr period (Sex *[F (1, 32) = 39.02, P< 0.001]*; AAV *[F (1, 32) = 6.49, P= 0.016])*. (I.) Depiction of viral infusion into the LH and representative image of expression. Quantification of viral mediated NOP deletion and heatmaps visualizing viral spread within the LH are shown in **Supplemental Fig. 4A-D**. (J.) Genetic deletion of NOP from the LH in Cre expressing mice resulted in blunted alcohol intake in females (Sex *[F (1, 30) = 48.56, P< 0.001]*; AAV *[F (1, 30) = 13.50, P< 0.001]*). (K.-L.) Sucrose and water intake were not affected by NOP deletion from the LH. However, females consumed more sucrose in 2-hr (Sex *[F (1, 30) = 27.86, P< 0.001]*) and more water over a 24-hr period (*Sex [F (1, 30) = 36.21, P< 0.001]*). (M) Depiction of viral infusion into the SuM and representative image of expression. Quantification of viral mediated NOP deletion and heatmaps visualizing viral spread within the SuM are shown in **Supplemental Fig. 5A-D**. (N.) Genetic deletion of NOP from the SuM resulted in a selective reduction of alcohol intake in female mice, with no effect in males (Sex *[F (1, 23) = 39.91, P< 0.001]*; AAV *[F (1, 23) = 7.48, P= 0.012]*). (O.) Sucrose intake was not affected by NOP deletion from the SuM. However, females consumed more sucrose in 2-hr (Sex *[F (1, 23) = 19.53, P< 0.001]*) (P.) Water intake over a 24-hr period was greater in female mice, and ANOVA revealed a factor interaction (Sex *[F (1, 23) = 15.38, P< 0.001]*; Sex*AAV *[F (1, 23) = 5.86, P= 0.024]*). However, post hoc analysis indicated no difference in water intake as a function of SuM NOP deletion. Data are represented as mean +/- SEM (*P< 0.05, **P< 0.01, ***P< 0.005, ****P< 0.001).

Because functional synapses were observed in LS^Pnoc^ terminal fields, genetic deletion studies were conducted to determine the role of NOP in the LS, LH, and SuM in relation to binge drinking behavior. AAV8-hSyn-GFP/cre was infused into the dLS of male and female NOP*fl/fl* mice (**Fig. 5E)**, which was largely confined to the dLS and iLS, and resulted in decreased *Oprl1* mRNA (**Supplemental Fig. 3A-D)**. After 4 weeks to allow for sufficient viral expression, alcohol drinking was assessed in the 2-hr modified DID paradigm for 3 weeks (**Fig. 5F)**. Females consumed more alcohol than males, and genetic deletion of NOP from the LS generally reduced alcohol drinking. However, this was primarily driven by males as there was no effect of knockdown in females. Average alcohol drinking across weeks is shown in **Supplemental Fig. 3E** and blood alcohol concentrations (BACs), alcohol intake, and BAC/intake correlation from the final day of drinking are shown in **Supplemental Fig. F-G**. The effect of LS-NOP deletion was selective to alcohol in males, but both sucrose drinking (**Fig. 5G**) and water intake (**Fig. 5H**) were attenuated in females. LS NOP deletion had no effect on locomotor activity or body weights (**Supplemental Fig. 3H-I**).

Genetic deletion of NOP from the LH was investigated in a separate study **Fig. 5I**. Infusion of a cre-expressing AAV into the LH was sufficient to decrease *Oprl1* mRNA and viral expression observed throughout the LH, centered on the tuberal LH with scant expression in neighboring ventromedial hypothalamus (**Supplemental Fig. 4A-D**). As in our previous studies, females consumed more alcohol than males and NOP deletion from the LH led to reduced alcohol intake in female, but not male mice (**Fig. 5J**). Alcohol intake by week, final day of drinking, BAC, and intake/BAC correlation can be found in **Supplemental Fig. 4E-G**. Sucrose and water intake were not affected by LH NOP deletion, but locomotor activity was attenuated in females (**Supplemental Fig. 4I**), suggesting the potential for a disruption of locomotion to affect alcohol drinking behavior.

Lastly, the effect of NOP deletion from the SuM on alcohol drinking was examined (**Fig. 5M**). Expression of cre in the SuM reduced *Oprl1* mRNA and viral expression was centered on the SuM, with some spread observed in the medial portions of the anterior VTA (**Supplemental Fig. 5 A-D**). Females consumed more alcohol than males and genetic deletion of NOP from the SuM resulted in a selective reduction of alcohol drinking in female mice (**Fig. 5N**). Drinking data by week, BACs on the final day of drinking and intake/BAC correlation can be found in **Supplemental Fig. 5 E-H**. SuM NOP deletion had no effect on sucrose (**Fig. 5O**) or water (**Fig. 5P**) intake, nor were any differences in locomotor activity observed (**Supplemental Fig. 5I**). While SuM NOP deletion was generally without effect on drinking males, body weight was reduced in cre-expressing males (**Supplemental Fig. 5J**).

## DISCUSSION

NOP antagonists are a promising treatment for AUD and rigorous examination of endogenous nociceptin circuitry is necessary to better understand the neurobiological basis for therapeutic efficacy. Here, we have identified a specific subpopulation of nociceptin-containing neurons within LS that shows functional changes following binge drinking, can bidirectionally modulate alcohol drinking behavior, and is activated during aversive experiences. We also show that the LS is likely an important site of action for NOP antagonists that reduce drinking in humans, and identify key hypothalamic projection sites involved in NOP modulation of alcohol drinking that will be important areas for future investigation.

### AUD Neurocircuitry: Role for the Lateral Septum in Binge Drinking and Withdrawal

The LS is a frequently overlooked node in the theoretical framework proposed for AUD, contributing to binge drinking and displaying responsivity during withdrawal. Repeated cycles of binge intoxication and withdrawal is proposed to generate an allosteric load, shifting the balance of excitatory/inhibitory systems that govern homeostasis in neuronal networks^53,54^. This imbalance ultimately biases circuit engagement that drives negative affect and excessive drinking behavior^49^. In line with these findings, LS^Pnoc^ neurons displays increased excitability during acute cessation of binge-like alcohol consumption at a timepoint when negative affective symptoms are observed^55^. Furthermore, withdrawal from alcohol produces a robust place aversion in rodents^56^ and here we show that optogenetic LS^Pnoc^ activation produces a similar effect. We speculate that alcohol-induced plasticity promotes hyperactivity of LS^Pnoc^ during withdrawal, generating a negative affective state that drives excessive drinking behavior. Indeed, the LS has been implicated in stress and anxiety states, and recent studies leveraging *in vivo* Ca imaging techniques have revealed heightened activity in genetically defined LS populations during exposure to stimuli associated with negative valence. For example, NTS-^46,57^, SST-^45^, and CRFR2-^44^ expressing neurons in the LS are responsive to footshock, and VGAT-expressing neurons are highly responsive to pain^58^. LS^Pnoc^ activity follows this pattern, displaying increased activity during footshock, air puff stimulus, sucrose spray, as well as exploration of the open arm in an EPM. However, LS^Pnoc^ activity is reduced during active grooming behavior when mice experience discomfort from a soiled coat. Silencing of hyperactive LS activity provoked by inflammatory pain has analgesic and anxiolytic properties^58^, suggesting that quiescence of LS activity provides relief from aversive stimuli. Indeed, alcohol consumption acutely suppresses activity of the dLS^59^ whereas acute withdrawal is associated with rebound excitation^60^. We speculate that LS^Pnoc^ activity/activation produces a negative internal state, mimicking aspects of withdrawal, and resultant excessive drinking behavior silences LS^Pnoc^ activity to provide relief from this state. This is an intriguing possibility and further in-depth study is warranted to address this possibility.

We have recently shown that activation of the dorsal septum (dSep; comprising the LS and dorsal tip of the medial septum) is sufficient to increase alcohol and sucrose intake in males, and increase locomotor activity^27^. Here we report that more discrete isolation of LS^Pnoc^ revealed a selective contribution to alcohol drinking, and the ability of this population to affect alcohol intake was not influenced by biological sex. Interestingly, LS^Pnoc^ Ca activity was not altered during active alcohol consumption and we speculate this was due to acute alcohol presentation, which was experimentally controlled as to not confound subsequent photometry experiments in the same mice. This resulted in relatively low levels of alcohol intake that are not associated with withdrawal or negative affect-driven drinking, which we speculate is necessary to uncover dynamic changes in LS^Pnoc^ activity.

The LS is heterogenous in terms of peptide expression and it is likely that targeting nociceptin-expressing neurons allowed for the uncovering of neuromodulation specific to alcohol. Regions governing binge drinking behavior are often associated with reward, and initial observations by Olds and Milner indicated that activation of the septal area is innately rewarding^61^. The LS ties directly into reward circuitry by innervating the VTA^62,63^, yet rodents will not self-stimulate for LS-VTA activation nor does activation of this pathway drive place preference^64^. Here, synaptophysin tracing experiments revealed a lack of LS^Pnoc^ innervation of the VTA, which agrees with retrograde tracing of VTA afferents in Pnoc-cre mice conducted by Parker and colleagues^65^. It is likely that any modulation of the VTA by LS^Pnoc^ is through indirect means, such as local microcircuits impinging upon non-Pnoc outputs to the VTA^39,42^, or modulation of downstream populations that project to the VTA. To this end, LS^Pnoc^ form monosynaptic inputs to the LH, which in turn innervates the VTA and regulates binge drinking, consummatory behaviors, and promotes aversive behaviors^66,67^. GABAergic neurons within the LH that project to the VTA directly promote consummatory behaviors^68–70^, while activation of glutamatergic neurons reduces feeding and reward-associated behaviors^71,72^. Given that LS^Pnoc^ form GABAergic synapses at the level of the LH, it is likely that activation of this circuit results in a suppression of LH-VGLUT neuronal activity allowing for the promotion of alcohol drinking behavior. The SuM is another major output of LS^Pnoc^ involved in arousal^73^, motivation^74^, and anxiety-like behavior^75^. Chemogenetic activation of the SuM is anxiolytic^75^, and silencing of LS^Pnoc^ may result in disinhibition thereby reducing alcohol-associated anxiety during withdrawal and subsequent motivation to consume alcohol. Consummatory behavior decreases SuM activity, and inhibition disrupts reward seeking but not consumption^76^.

### NOP as Target for AUD Pharmacotherapy

The ability of NOP antagonists to reduce alcohol in rodent models is well documented, where treatment has reduced binge-like alcohol consumption in male mice^21^, as well as attenuate drinking and reinstatement of alcohol-seeking in rats^18–20^. A few discrete sites of action have been explored, indicating that blockade of NOP in the VTA or CeA, but not NAc, decreased excessive alcohol drinking in male rats^19,20^. Here we report that NOP in LS, LH, and SuM also regulate excessive drinking and argue for the exploration of antagonist effects outside of defined mesolimbic and extended amygdala addiction circuity, and this is consistent with global genetic NOP knockout which suppresses binge drinking^21^ and alcohol self-administration^77^. Moreover, our findings suggest that the ability of NOP to regulate alcohol consumption was dependent upon sex, and inclusion of sex as a biological variable (SABV) is critical to forming the preclinical basis for the clinical pursuit of system targets in the treatment of AUD. The present studies indicate that the ability of a NOP antagonist to reduce alcohol drinking may be driven by different sites of NOP expression in the male and female brain linked to different motivations to consume alcohol. Sex differences exist in the motivation to consume alcohol, where initial excessive drinking in men is reported to be driven primarily by positive reinforcement mechanisms^78^ whereas, women tend to initiate excessive drinking later in life and rapidly escalate in a telescoping effect that is thought to be driven primarily by negative reinforcement^79,80^. Genetic deletion of NOP from the LS selectively reduced alcohol intake in male mice, without affecting alcohol intake in females, suggesting a link to positive reinforcement. However, LS NOP deletion attenuated both sucrose and water intake in females, indicating a fundamental difference in motivational processing between sexes. In contrast, NOP deletion from the LH attenuated alcohol drinking in females without affecting sucrose or water intake, suggesting a selective reduction in the motivation to consume alcohol. However, this was accompanied by a nonspecific attenuation of locomotor activity, making interpretation difficult. A more clear story emerges at the level of the SuM, where NOP deletion decreased alcohol intake in females with no effect on sucrose intake, water intake, or locomotor activity. We speculate that deletion of NOP results in disinhibition of the SuM that decreases the motivation to consume alcohol. This hypothesis is supported by recent studies indicating that chemogenetic activation of the SuM is anxiolytic, and decreased anxiety is associated with decreased alcohol consumption^75^.

Sex-specific effects were also observed in relation to LS^Pnoc^ excitability, where cells from females were largely insensitive to a history of binge-like alcohol consumption. It is important to note that 3 weeks of binge drinking in the DID model is a relatively brief window of voluntary alcohol drinking, and more robust models of alcohol drinking and/or exposure, such as the chronic intermittent ethanol vapor model, may drive adaptations in LS^Pnoc^ in females that do not emerge in the DID model. Notably, we did not assess neurotransmitter release dynamics of LS^Pnoc^ neurons, which could be a potential site of action that is altered following alcohol consumption, and may underlie sex-differences at the synaptic level either within the LS itself or in downstream projection sites. However, chemogenetic silencing/activation bidirectionally modulated alcohol intake independent of sex, suggesting that LS^Pnoc^ manipulation is sufficient to affect voluntary alcohol intake regardless of sex differences in sensitivity to alcohol.

In conclusion, the present studies identify LS^Pnoc^ as a GABAergic population that is highly expressed in the dLS. These neurons are responsive to stimuli associated with negative valence and acute activation has aversive qualities. Furthermore, these neurons display increased excitability during withdrawal from binge-like alcohol consumption and play a causal role in alcohol drinking, in that chemogenetic activation increased alcohol drinking, whereas inactivation decreased alcohol intake. The effect of LS^Pnoc^ manipulation to influence alcohol drinking was not due to a general shift in consumption of a palatable fluid as sucrose drinking was unaffected, nor was this due to nonspecific locomotor effects. LS^Pnoc^ efferent tracing revealed terminal fields within the LS itself, as well as the LH and SuM and these were confirmed to be monosynaptic GABAergic projections. Lastly, NOP were found to contribute to alcohol drinking in a sex-dependent fashion in that NOP deletion from the LS decreased alcohol intake in males, whereas NOP deletion from the LH and SuM decreased alcohol drinking in females. Together, these findings identify septohypothalamic nociceptin signaling as a previously unidentified system regulating excessive alcohol consumption.

---

**Supplemental Fig. 1.**
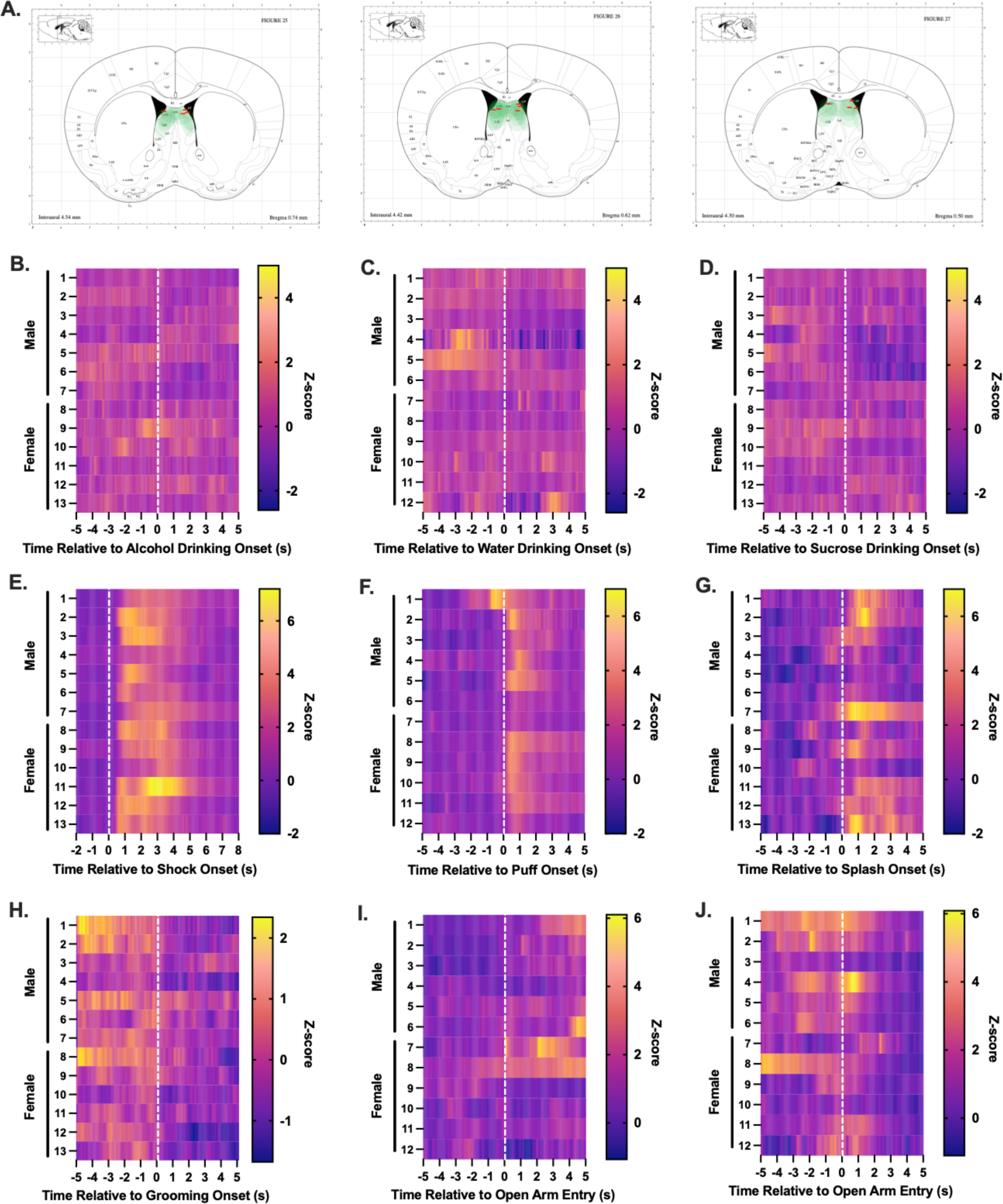
(A.) Representative heatmap of viral expression and fiber placements in Pnoc-cre mice receiving AAV-hSyn-DIO-GCaMP7f infusion into the LS. Fiber tip placements are depicted as a red line. (B.-J.) Peri-event heatmaps for individual subject LS^Pnoc^ GCaMP activity during (B.) water drinking, (C.) alcohol drinking, (D.) sucrose drinking, (E.) footshock exposure, (F.) air puff exposure, (G.) sucrose splash exposure, (H.) grooming post sucrose splash, (I.) open arm entry in EPM, and (J.) open arm exit in EPM. Data are represented as mean.

**Supplemental Fig. 2.**
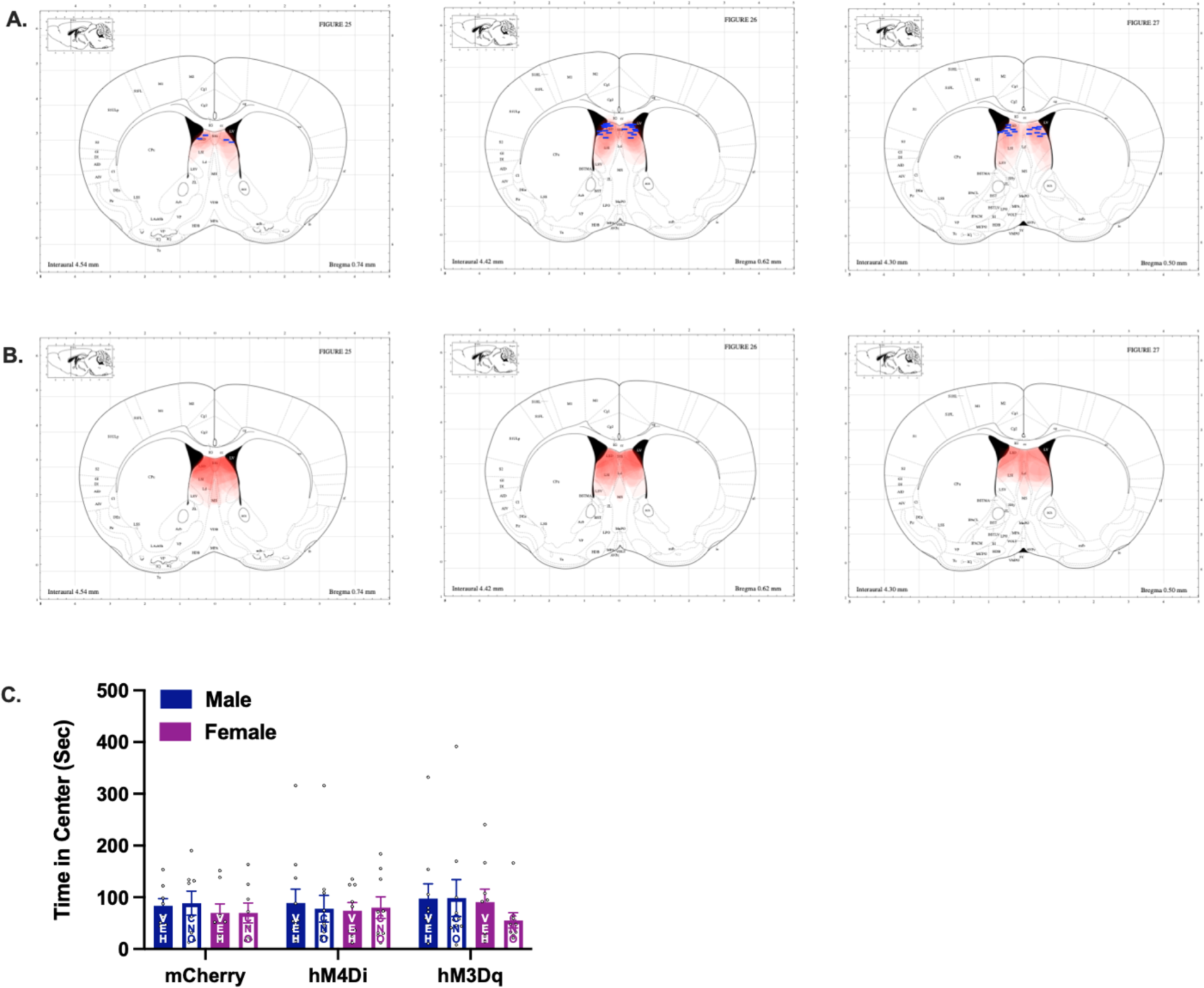
(A.) Representative heatmap of viral expression and optical fiber placements in Pnoc-cre mice receiving AAV8-hSyn-DIO-ChR2/mCherry infusion into the LS. Fiber tip placements are depicted as a blue line. (B.) Representative heatmap of mCherry expression in Pnoc-cre mice receiving AAV-hSyn-DIO-hM3Dq/hM4di/mCherry infusion into the LS. (C.) Chemogenetic manipulation of LS^Pnoc^ did not affect time spent in the center of an open field in male and female mice. Data are represented as mean +/- SEM (*P< 0.05, **P< 0.01, ***P< 0.005, ****P< 0.001).

**Supplemental Fig. 3:**
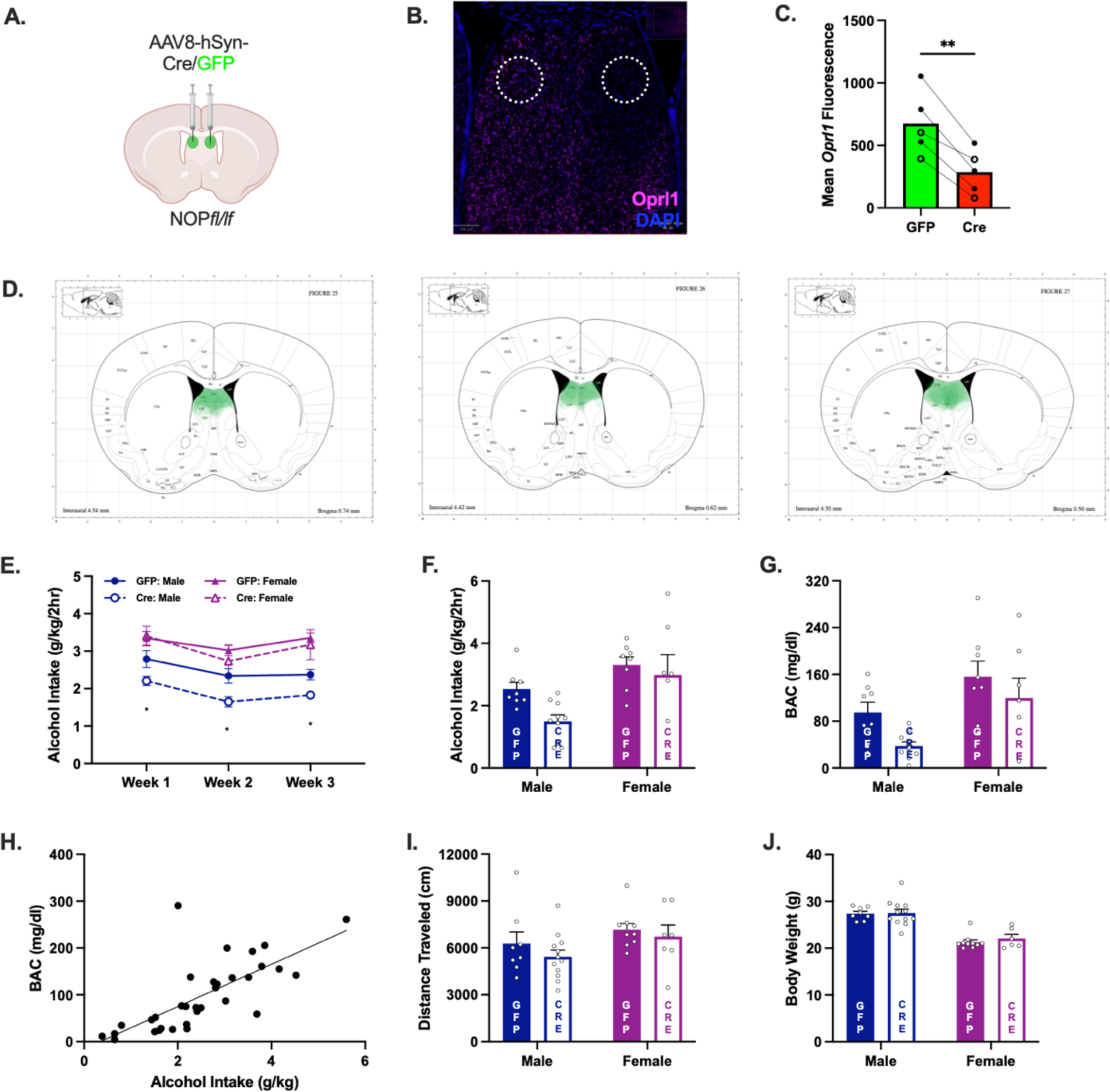
NOP deletion from the LS. (A.) Representative image of viral infusion strategy for NOP genetic deletion in the LS. (B.) Representative *in situ* hybridization image of *Oprl1* mRNA in mice with unilateral AAV8-hSyn-GFP-cre into the LS. (C.) Quantification of *Oprl1* immunofluorescence in NOP*fl/fl* mice in the LS. Unilateral infusion of Cre-expressing AAV resulted in decreased *Oprl1* mean fluorescence compared to the control side (Paired t-test; *t = 6.621, df = 4; P= 0.003*). Males = closed circle, Females = open circle. (D.) Representative heatmaps depicting AAV8-hSyn-GFP/cre expression within the LS of NOP*fl/fl* mice. (E.) Average weekly alcohol intake. Genetic deletion of NOP from the LS decreased alcohol intake in male mice across weeks (AAV: *[F (1, 32) = 5.65, P= 0.024]*; Time *[F (1, 32) = 5.08, P= 0.031]*), and females generally consumed more alcohol than males (Sex*: [F (1, 32) = 38.77, P< 0.001]*). (F.) Alcohol intake on the final day of drinking prior to blood collection. Females consumed more alcohol than males (Sex: *[F (1, 29) = 10.71, P= 0.003]*) and the ability of LS NOP deletion to affect alcohol drinking in males neared significance (P= 0.066). (G.) Blood alcohol concentration after the final day of drinking. Deletion of NOP from the LS had a modest effect on reducing resultant BAC post alcohol drinking (Sex: *[F (1, 29) = 10.95, P= 0.003]*; AAV: *[F (1, 29) = 4.72, P= 0.038]*). (H.) Relationship between alcohol consumed and BAC. The amount of alcohol consumed during the 2-hr drinking session was positively associated with BAC (Linear regression: *R2 = 0.518, F = 33.36, P< 0.001*). (I.) Distance traveled in an open field. LS NOP deletion did not affect locomotor activity. (J.) Body weight. NOP deletion did not affect body weight, but males generally had greater weight than females (Sex: *[F (1, 32) = 63.27, P< 0.001]*). Data are represented as mean +/- SEM (*P< 0.05, **P< 0.01, ***P< 0.005, ****P< 0.001).

**Supplemental Fig. 4:**
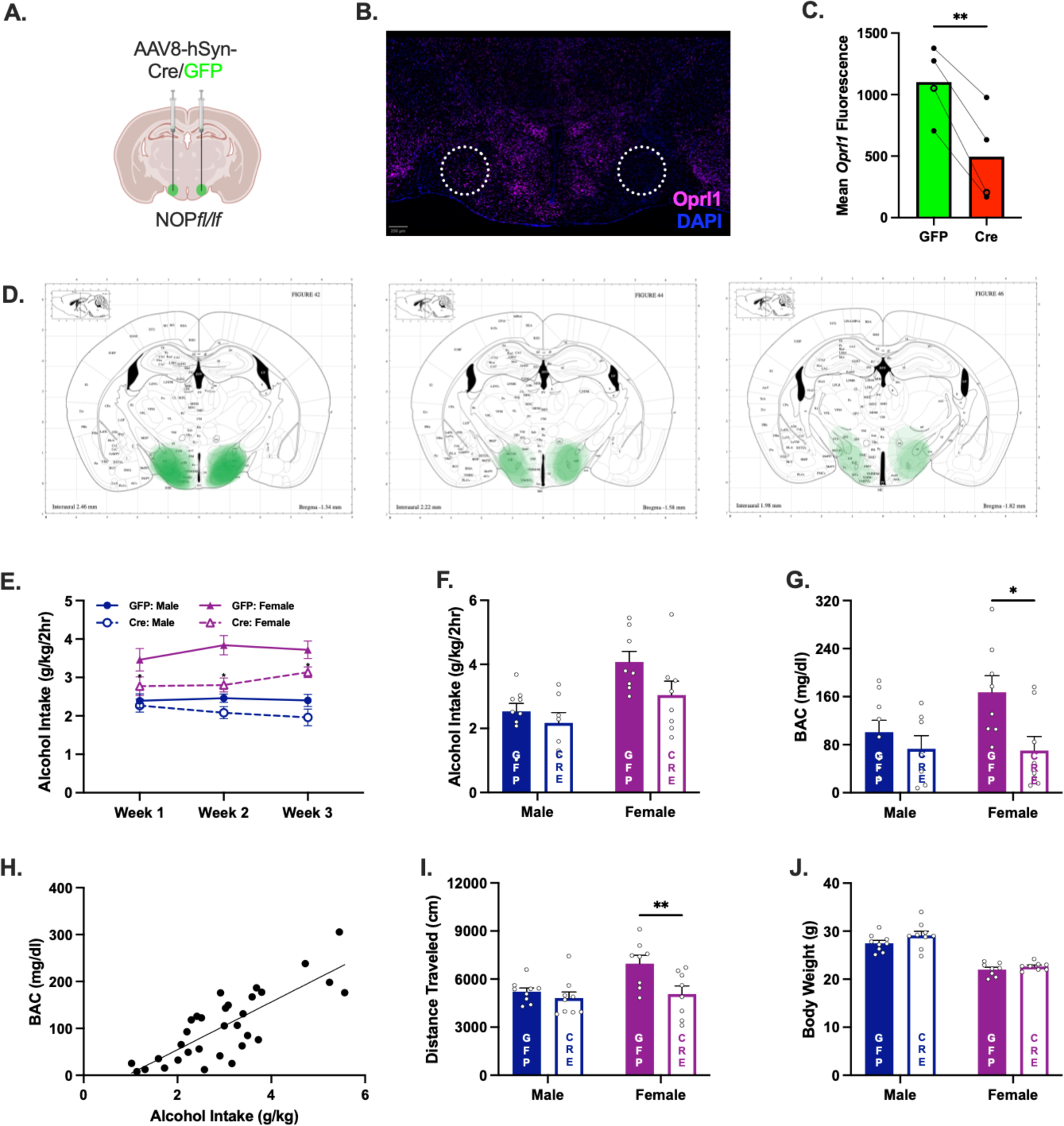
NOP deletion from the LH. (A.) Representative image of viral infusion strategy for NOP genetic deletion in the LH. (B.) Representative *in situ* hybridization image of *Oprl1* mRNA in mice with unilateral AAV8-hSyn-GFP-cre into the LH. (C.) Quantification of *Oprl1* immunofluorescence in NOP*fl/fl* mice in the LH. Unilateral infusion of Cre-expressing AAV resulted in decreased *Oprl1* mean fluorescence compared to the control side (Paired t-test; *t = 6.407, df = 3, P= 0.008*). Males = closed circle, Females = open circle. (D.) Representative heatmaps depicting AAV8-hSyn-GFP/cre expression within the LH of NOP*fl/fl* mice. (E.) Average weekly alcohol intake. Genetic deletion of NOP from the LH decreased alcohol intake in female mice across weeks (AAV: *[F (1, 30) = 13.50, P< 0.001]*), and females generally consumed more alcohol than males (Sex: *[F (1, 30) = 48.56, P< 0.001]*). (F.) Alcohol intake on the final day of drinking prior to blood collection. Females generally consumed more alcohol than males (Sex [*F (1, 28) = 12.60, P= 0.001*]; AAV [*F (1, 28) = 4.215, P= 0.0495*]) and the ability if NOP deletion to affect alcohol intake neared significance in females (P= 0.078). (G.) Blood alcohol concentration after the final day of drinking. BAC was attenuated in females lacking NOP from the LH compared to controls (AAV [*F (1, 28) = 7.14, P= 0.012*]). (H.) Relationship between alcohol consumed and BAC. The amount of alcohol consumed during the 2-hr drinking session was positively associated with BAC (Linear Regression: *R2 = 0.638, F = 52.83, P< 0.001*). (I.) Distance traveled in an open field. Sex [*F (1, 30) = 5.82, P= 0.022*]; AAV [*F (1, 30) = 7.68, P= 0.010*]. (J.) Body weight. NOP deletion did not affect body weight, but males generally had greater weight than females (Sex: *[F (1, 30) = 91.18, P< 0.001]*). Data are represented as mean +/- SEM (*P< 0.05, **P< 0.01, ***P< 0.005, ****P< 0.001).

**Supplemental Fig. 5:**
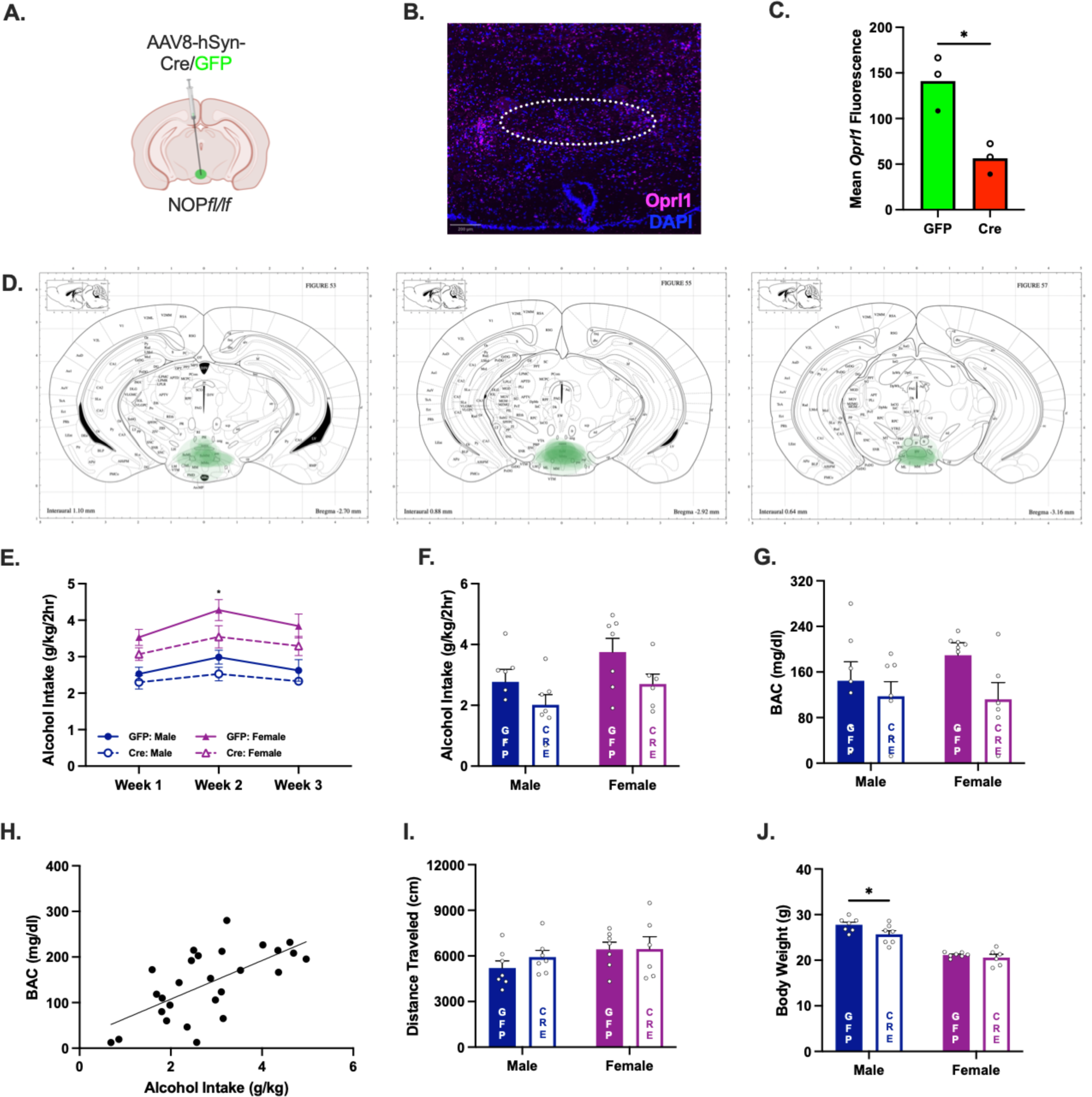
NOP deletion from the SuM. (A.) Representative image of viral infusion strategy for NOP genetic deletion in the SuM. (B.) Representative *in situ* hybridization image of *Oprl1* mRNA in mice with unilateral AAV8-hSyn-GFP-cre into the Sum. (C.) Quantification of *Oprl1* immunofluorescence in NOP*fl/fl* mice in the SuM. Unilateral infusion of Cre-expressing AAV resulted in decreased *Oprl1* mean fluorescence compared to control mice (Unaired t-test; *t = 4.302, df = 4, P= 0.013*). Males = closed circle, Females = open circle. (D.) Representative heatmaps depicting AAV8-hSyn-GFP/cre expression within the SuM of NOP*fl/fl* mice. (E.) Average weekly alcohol intake. Females generally consumed more alcohol than males post hoc analysis revealed a reduction in alcohol drinking in female mice during the second test week after SuM NOP deletion (Sex [*F (1, 23) = 40.02, P< 0.001*]; AAV [*F (1, 23) = 7.58, P= 0.011*]). (F.) Alcohol intake on the final day of drinking prior to blood collection. Females generally consumed more alcohol than males and deletion of NOP from the SuM had a general effect of reducing alcohol intake (Sex [*F (1, 23) = 4.56, P= 0.044*]; AAV [*F (1, 23) = 5.32, P= 0.031*]). (G.) Blood alcohol concentration after the final day of drinking. Genetic deletion of NOP from the SuM did not affect resultant BACs on the final day of alcohol drinking (AAV [F (1, 23) = 3.530 P= 0.073]). (H.) Relationship between alcohol consumed and BAC. The amount of alcohol consumed during the 2-hr drinking session was positively associated with BAC (Linear Regression: *R2 = 0.423, F = 18.29, P< 0.001*). (I.) Distance traveled in an open field. Cumulative distance traveled was not affected by deletion of NOP from the SuM. (J.) Body weight. While males generally had greater body weights than females, NOP deletion from the SuM resulted in decreased body weight in male mice (Sex [*F (1, 23) = 91.26, P< 0.001*]; AAV [*F (1, 23) = 4.80, P= 0.039*]). Data are represented as mean +/- SEM (*P< 0.05, **P< 0.01, ***P< 0.005, ****P< 0.001).

**Supplemental Table 1:**
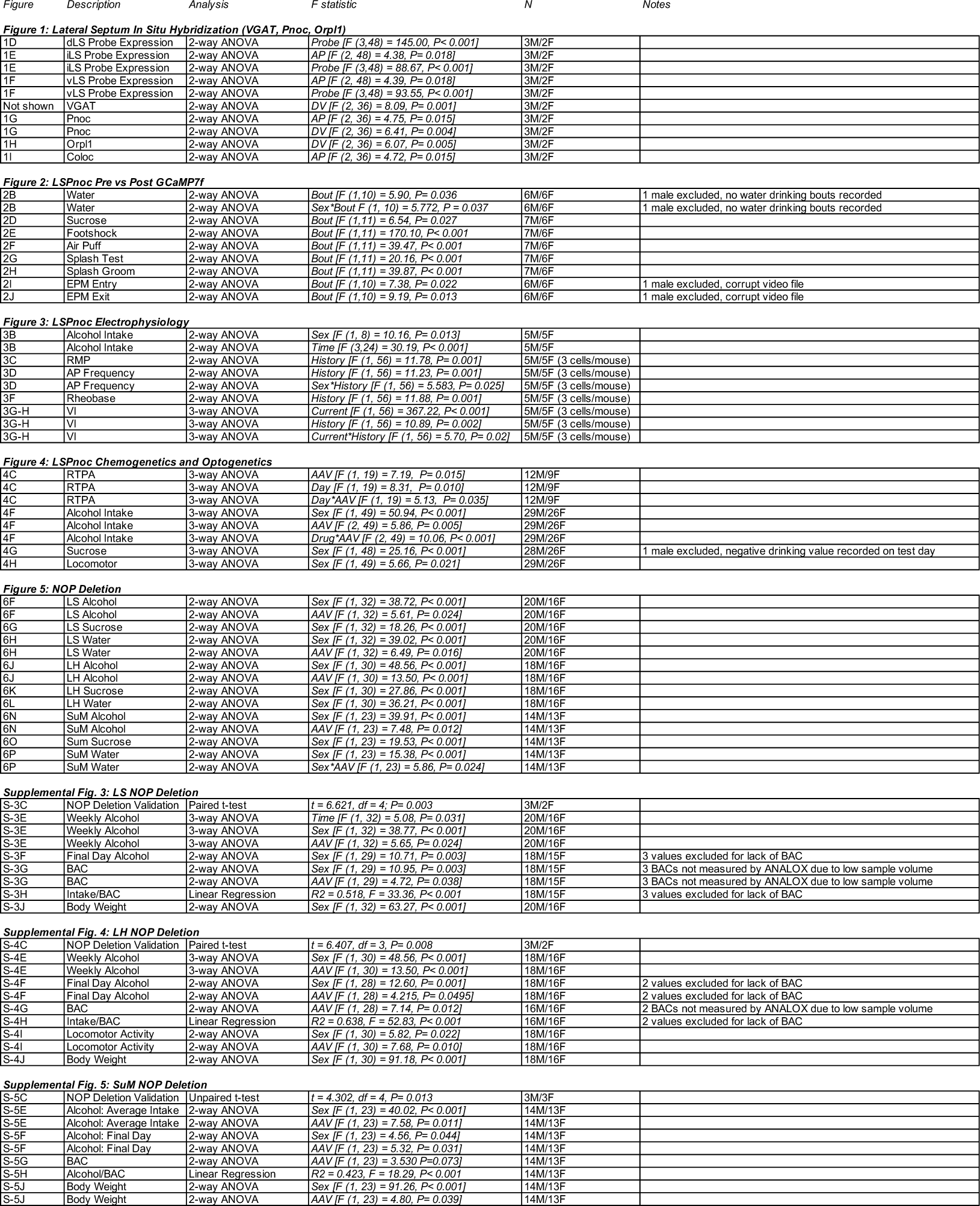
Statistics.

